# Modeling protein-small molecule conformational ensembles with PLACER

**DOI:** 10.1101/2024.09.25.614868

**Authors:** Ivan Anishchenko, Yakov Kipnis, Indrek Kalvet, Guangfeng Zhou, Rohith Krishna, Samuel J. Pellock, Anna Lauko, Gyu Rie Lee, Linna An, Justas Dauparas, Frank DiMaio, David Baker

**Affiliations:** Department of Biochemistry, University of Washington, Seattle, WA 98105, USA; Institute for Protein Design, University of Washington, Seattle, WA 98105, USA; Graduate Program in Biological Physics, Structure and Design, University of Washington, Seattle, WA 98105, USA; Howard Hughes Medical Institute, University of Washington, Seattle, WA 98105, USA

## Abstract

Modeling the conformational heterogeneity of protein-small molecule interactions is important for understanding natural systems and evaluating designed systems, but remains an outstanding challenge. We reasoned that while residue level descriptions of biomolecules are efficient for de novo structure prediction, for probing heterogeneity of interactions with small molecules in the folded state an entirely atomic level description could have advantages in speed and generality. We developed a graph neural network called PLACER (Protein-Ligand Atomistic Conformational Ensemble Resolver) trained to recapitulate correct atomic positions from partially corrupted input structures from the Cambridge Structural Database and the Protein Data Bank; the nodes of the graph are the atoms in the system. PLACER accurately generates structures of diverse organic small molecules given knowledge of their atom composition and bonding, and given a description of the larger protein context, builds up structures of small molecules and protein side chains for protein-small molecule docking. Because PLACER is rapid and stochastic, ensembles of predictions can be readily generated to map conformational heterogeneity. In enzyme design efforts described here and elsewhere, we find that using PLACER to assess the accuracy and pre-organization of the designed active sites results in higher success rates and higher activities; we obtain a preorganized retroaldolase with a *k*_cat_/*K*_M_ of 11000 M^-1^min^-1^, considerably higher than any pre-deep learning design for this reaction. We anticipate that PLACER will be widely useful for rapidly generating conformational ensembles of small molecule and small molecule-protein systems, and for designing higher activity preorganized enzymes.

## Main Text

Interactions of proteins with nucleic acids, small organic and inorganic molecules, and metals are critical to biological function, but atomistic modeling of such interactions and their conformational heterogeneity remains a challenging problem. Deep learning (DL)-based small molecule docking tools like DiffDock(1) improve on earlier methods such as Glide(2) and GNINA/SMINA, but in the high accuracy regime the difference in performance is not pronounced, and performance also drops substantially on unseen receptors(1, 3, 4). DL methods have also been devised to generate small molecule conformations from their chemical structure(5–7); when trained on the synthetic GEOM-DRUGS dataset(8) of drug-like molecules computed with semi-empirical QM methods, diffusion generative models like DiffDock(1) and Torsional Diffusion(7) show best-in-class performance on the hold-out test set. However, these approaches model specific classes of interaction partners, limiting the ability to model the full spectrum of protein functions, and the input features can differ depending on the type of input molecules, which could limit the ability of the networks to distill general physico-chemical principles. AlphaFold2 (AF2)(9) and RoseTTAFold (RF)(10) enabled atomically accurate structure prediction of proteins and protein-protein complexes using sequences and structures of evolutionary-related proteins as inputs. These methods have recently been extended by AF3 and related approaches to modeling the structures of protein-nucleic acid (NA) complexes and more general biomolecular systems(11–13) by supplementing tokens representing residues with full atomic representations of the non-protein components of the system.

We reasoned that rapid and accurate modeling of structure and conformational heterogeneity could be achieved by an atom centric approach to predicting the structures of small molecules and peptides in isolation and in the context of a protein binding or active site defined at the backbone level. Networks such as AF2, AF3, and RF achieve accurate structure prediction using a primarily residue-level description of biomolecules and multiple sequence information to help infer contacts. At the atomic level close to the native structure, the complexity grows considerably and evolutionary information is less relevant. Hence, we reasoned that an appropriate starting point for an ensemble generation method would not be the amino acid sequence of a protein but rather the coordinates of the overall protein backbone surrounding a binding or catalytic pocket, along with an atomic level description of the bonded geometry of the interacting small molecules and amino acid side chains. Such a network would, by construction, not be useful for structure prediction from sequence, but could be very useful for modeling the conformations of small molecules and constrained peptides both in isolation and in a protein binding site along with the conformations of the interacting sidechains. Since the protein backbone structure is input, the calculations could be faster than AF2, AF3, and RF, enabling rapid binding site refinement and evaluation. We reasoned that a network that generated predictions stochastically could have the advantage of enabling rapid generation of predicted ensembles for modeling systems with a distribution of possible states (which is difficult with AF2, AF3, and RF), and for evaluating the extent of preorganization of designed molecules and functional sites.

Guided by this reasoning we set out to develop a stochastic deep neural network called PLACER (Protein-Ligand Atomistic Conformational Ensemble Resolver) for atomistic modeling of small molecules and small molecule-protein interactions (Fig. 1A). In many applications, a reliable structure of the target protein can be generated by AF2(9) or RF(10), or obtained from the PDB, and location of the putative interacting region on the protein surface is also known; the task is then to dock a small molecule into the region of interest, while adjusting conformations of the protein side chains and the molecule itself. We framed the learning problem as a structure denoising task and trained PLACER to recapitulate the correct atom positions from partially corrupted input structures provided that all the chemical information about the system being modeled is known from the start. We customized the corruption strategy to the application of interest. In the case of the protein-small molecule docking (Fig. 1A), the inputs to the network include the protein backbone coordinates, the amino acid sequence with side chain coordinates randomly initialized around the respective C-alpha atoms, and the small molecule chemical structure with atomic positions randomly initialized in the vicinity of the putative binding site.

**Figure 1.**
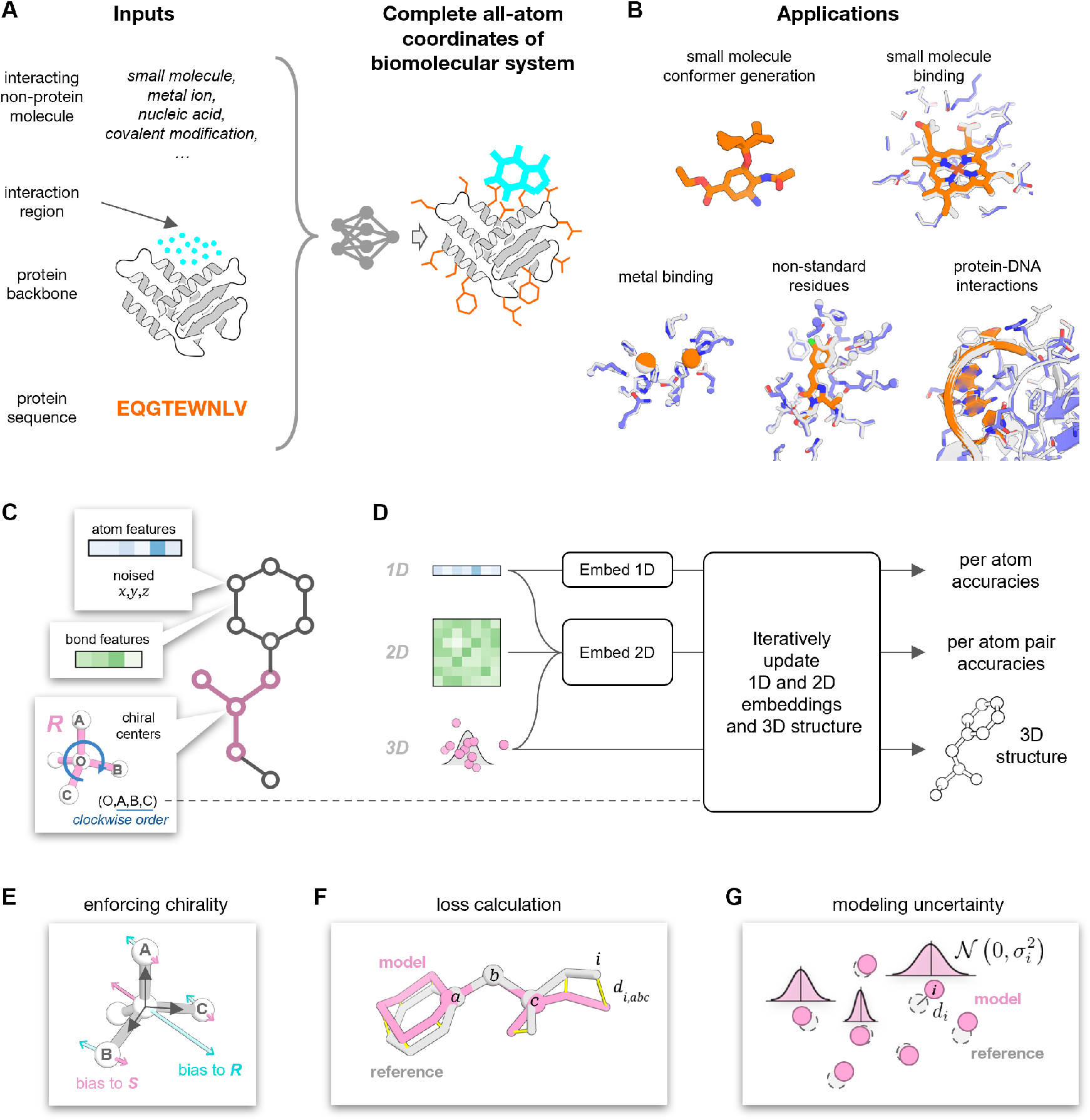
Overview of PLACER. **A**) PLACER is a denoising neural network which takes at input a partially corrupted protein structure and the chemical structure (but not the coordinates) of any interacting molecules, and predicts the all-atom structure of the complex, as well as the uncertainties in the atom positions in the generated model. **B**) PLACER can be used for a wide range of tasks including docking of small molecules and metals to a protein target, modeling non-standard residues, and predicting side chain conformations of amino acid residues and nucleotides at the protein-DNA interface. Shown are x-ray structures (in gray) superimposed with PLACER models (in blue and orange). **C**) At input, the molecular system is represented by an annotated graph where nodes are individual atoms and edges are chemical bonds between atoms. Information about chiral centers is supplied to the network as (O,A,B,C) tuples where O is the central pyramidal or tetrahedral atom and its neighboring atoms A,B,C are ordered clockwise. **D**) PLACER is a three-track network that iteratively updates 1D and 2D embeddings and the 3D structure, producing at each iteration a refined atomic structure model and estimating uncertainties in atom placements. **E**) The triple product 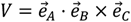 of the three unit vectors pointing from the central chiral atom to its neighbors (gray arrows) is a pseudoscalar that differs in sign for the *R* and *S* configurations: for ideal tetrahedral geometry 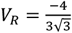 and 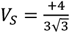. By comparing *V* in the non-ideal geometry of the modeled structure to the ideal values *V*_*R*_ or *V*_*S*_ and taking gradients w.r.t. atom coordinates, one gets biasing vectors 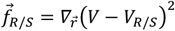 showing the directions in which atoms should be moved around in order to recreate the desired configuration. **F**) All-atom *FAPE* is calculated by aligning the model and the reference structures on every three respective bonded atoms *a, b, c* and calculating the deviations in atom positions between the aligned structures. *FAPE*_*all atom*_ is then the mean over all atoms and all superimpositions. Atom-atom distances are clamped at 10Å. **G**) Assuming that uncertainties in atom positions in the modeled structure are normally distributed, we let the dedicated head of the network predict the variances *σ*_*i*_^2^ for every atom in the system to recapitulate actual deviations *d*_*i*_. These variances are learned during training by maximizing the likelihood 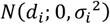.

At input, the molecular system is converted to a chemical graph with nodes representing individual heavy atoms (hydrogen atoms are not modeled to reduce computation cost) and edges representing chemical bonds between atoms (Fig. 1C). This representation is uniform across molecule types. Each node in the network carries information about the atom type and its 3D coordinates which are initially corrupted. The network is tasked to iteratively denoise the input coordinates and to estimate uncertainties in the atom positions of the output model structure (Fig. 1D). PLACER has a three-track architecture inspired by RF (10). After the initial embedding of the 1D and 2D features, they are passed to a block which iteratively updates the embeddings and the 3D structure. In the iteration block, the atom neighbor graph is first constructed: for every atom, the 32 closest neighbors are picked in equal proportions based on both spatial proximity and proximity in the chemical graph. 2D pair features are then projected by a feed-forward adapter layer to edge embeddings, and, together with the 1D features, the atom neighbor graph and the current atomic 3D structure serve as inputs to the SE3-Transformer network(14) which updates the 3D coordinates and the 1D embeddings. Information about the chiral centers is communicated to the network via *type-1* (vector) features (Fig. 1E). Features in the 2D track undergo pair-to-pair updates with bias from structure(15). Atom and atom pair confidence prediction heads branch off the 1D and 2D tracks respectively, completing the iteration block. The fully trained network consists of eight iteration blocks with shared weights.

PLACER is trained using a combination of structure and confidence prediction losses applied after every iteration. The primary structure loss is all-atom frame-aligned point error *FAPE*_*all atom*_, which is an extension of the original *FAPE* from Ref (9) to arbitrary molecules (Fig. 1F). The confidence of the modeled structures is evaluated on a per atom and per atom pair basis. Atom-atom pair accuracies are estimated using distance signed error approach introduced in Ref (16). On the per-atom level, we predict all-atom lDDT scores as in AF2 (9). The network also predicts deviations in atomic positions *σ*_*i*_ (in Å) in the generated model relative to the reference structure (Fig. 1G). These predicted deviations can then be combined over a subset of atoms of interest (e.g., a small molecule) to give the predicted RMSD 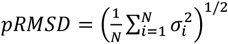.

### Tuning network architecture on experimental structures of small molecules

While small rigid molecules have known three-dimensional structures, larger organic molecules can have unique ground states that are very time consuming to predict with computational methods, hence, accurate prediction of general molecular structure is an important challenge. We explored network architecture and training hyperparameters on crystal structures of chemicals from the Cambridge Structural Database(17). This also helped ensure that the network architecture can handle diverse chemistry and considerably reduced the architecture exploration time (PLACER training on the PDB until convergence takes x10 longer). The training task was to predict the small molecule conformation observed in the crystal from the chemical structure and randomly initialized starting coordinates (Fig. 2A). The training and validation datasets consisted of 226,684 and 7,116 examples of organic non-polymeric small molecules, respectively (further details can be found in the SI, section 1.2), and confidence heads were omitted. As PLACER is a denoising network, structurally diverse samples of molecular conformations can be generated by running the network multiple times with different random initialization of the input coordinates. Training of PLACER on CSD structures was done in two stages: in the first stage, four iteration blocks were used and *FAPE*_*all atom*_ was the only loss term. In the second stage, the number of iterations was increased to eight and bonded geometry loss terms were added to enhance the local quality of predicted structures (red vs violet on Figs. 2B and C). Reducing the number of iterations to two or replacing the *FAPE* loss with a combination of coordinate and distance RMSD losses both result in predictions of substantially lower quality (orange and blue on Figs. 2B and C). Performance also degraded when the 2D inputs did not include the bond separation feature (an integer counting the number of covalent bonds between any two atoms in the chemical graph) but only included a binary feature flagging that the two atoms are immediately connected by a bond. Fully trained PLACER generates the correct 3D structures of molecules as complex as macrocycles with over 50 atoms (Fig. 2D) with sub-Å accuracy, including peptidic macrocycles(18) shown in the top row of Fig. 2D.

**Figure 2.**
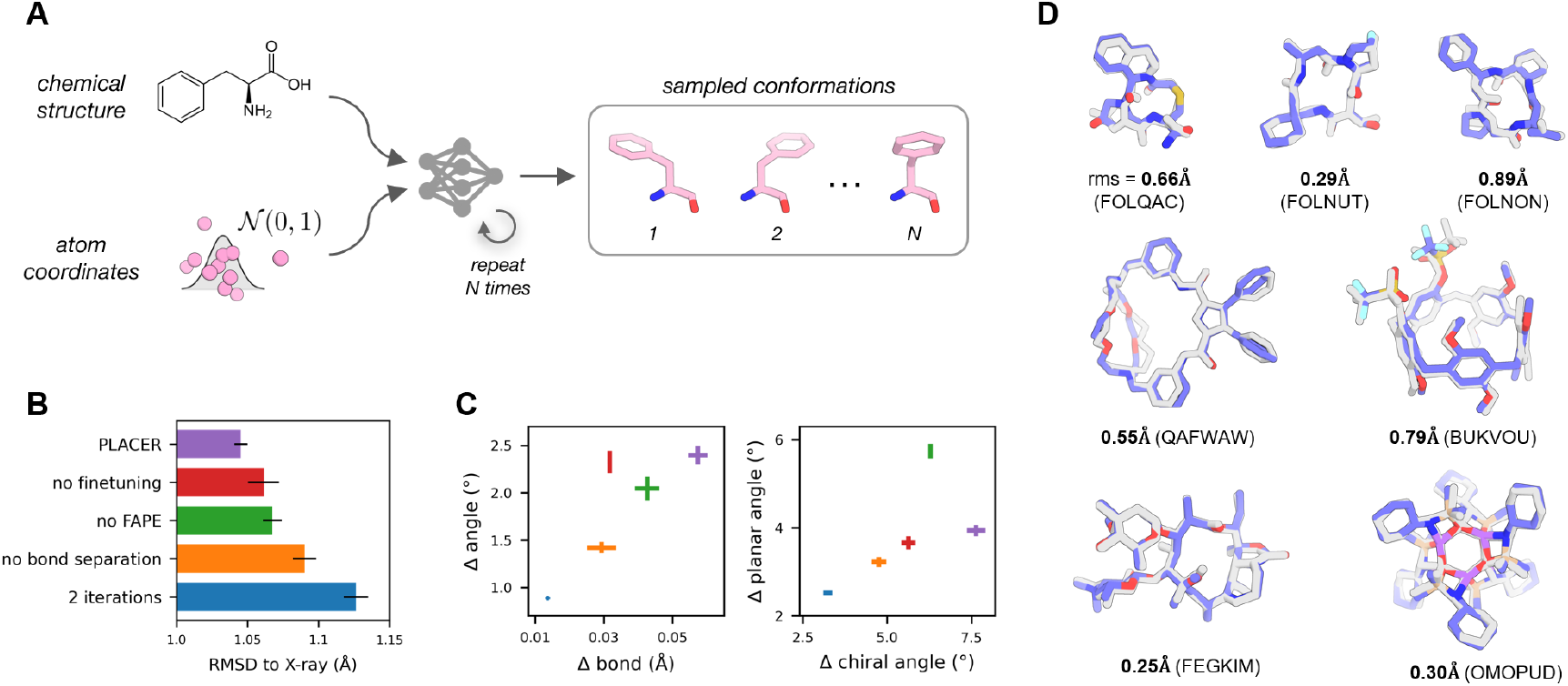
Modeling complex small molecules. **A)** PLACER predicts the 3D structure of a small molecule from its chemical structure and randomly initialized atomic coordinates. Running PLACER multiple times with different random initializations of atom positions yields a diverse set of molecular conformations. **B)** Ablating PLACER features negatively affects performance as measured by the root-mean-square deviation in atom coordinates between the model and the reference x-ray structure after the two are optimally superimposed. **C)** Quality of the local geometry in models produced by the five networks from panel B. Shown are mean absolute errors in bonds, bonded, planar, and chiral angles computed from the model and reference structures. Validation set of 7,116 small molecules from the CSD was used for panels B and C. A single conformer was generated for every validation example; the error bars represent standard deviations over 5 independent validation runs. **D)** Examples of seven macrocyclic molecules(18) for which PLACER generates the correct structure. Shown are the best RMSD models (in blue; out of 1000 generated) overlaid with the experimental crystal structures (gray). CCDC database identifiers shown in parentheses.

### Modeling protein-small molecule interactions with PLACER

To train PLACER on the structures from the PDB (proteins plus small molecules, rather than small molecules alone, as in the case of the CSD) we parsed them into chemical graphs (Fig. 1C) using the Chemical Component Dictionary(19) that provides detailed chemical description for all residue and small molecule components found in the PDB. We reasoned that although some molecules in the PDB may be non-biological (e.g., solvents) and their interactions with the rest of the structure may be non-specific, these interactions could still carry information about physico-chemical preferences at molecular interfaces and thus be informative for the network training; only water molecules were excluded. Training and validation sets (112,828 and 7,090 examples, respectively) were compiled from higher resolution (<2.5Å) structures deposited to the PDB before January 12, 2023. During training, the input structures were cropped to at most 600 heavy atoms, and centered around a randomly selected atom. The non-backbone-atom coordinates of all individual connected molecular components in the crop were collapsed to their connected backbone atom (if residue or nucleic acid), or to a randomly selected atom (ligands), and corrupted with Gaussian noise (*σ* = 1.5Å). From these starting points, PLACER was trained to recapitulate the atomic coordinates of all atoms in the cropped regions. For a detailed description of the training procedure, see Supplementary Methods.

We used PLACER to generate ensembles of small molecule poses in the pocket of the target protein (Fig. 3A) by repeated runs with different random initialization of the input coordinates. For each run, we chose one ligand atom at random, added Gaussian noise (σ = 1.5 Å) to its coordinates, collapsed all other ligand atoms onto that atom (these starting positions from multiple network runs are shown in Fig. 2B and D), and added uncorrelated Gaussian noise (σ = 1.5 Å) to the resulting coordinates of all ligand atoms — exactly matching the initialization used during training and ensuring that neither the internal structure of the ligand nor its orientation in the pocket were inferrable from the inputs to the network. Analyzing the resulting ensembles reveals that PLACER is not very sensitive to the initial placement of the ligand: multiple different starting positions yield near native predictions (Fig. 3B), and these positions cover the entire space sampled at input (Fig. 3D). We also observed that the predicted *pRMSD* score calculated over the ligand atoms allows selection of more accurate models from the sampled pool (Fig. 3E), with the best scoring models closely matching the experimental structure (ligand RMSD = 0.53Å and 0.77Å for heme(20) and cortisol(21) respectively, Fig. 3C).

**Figure 3.**
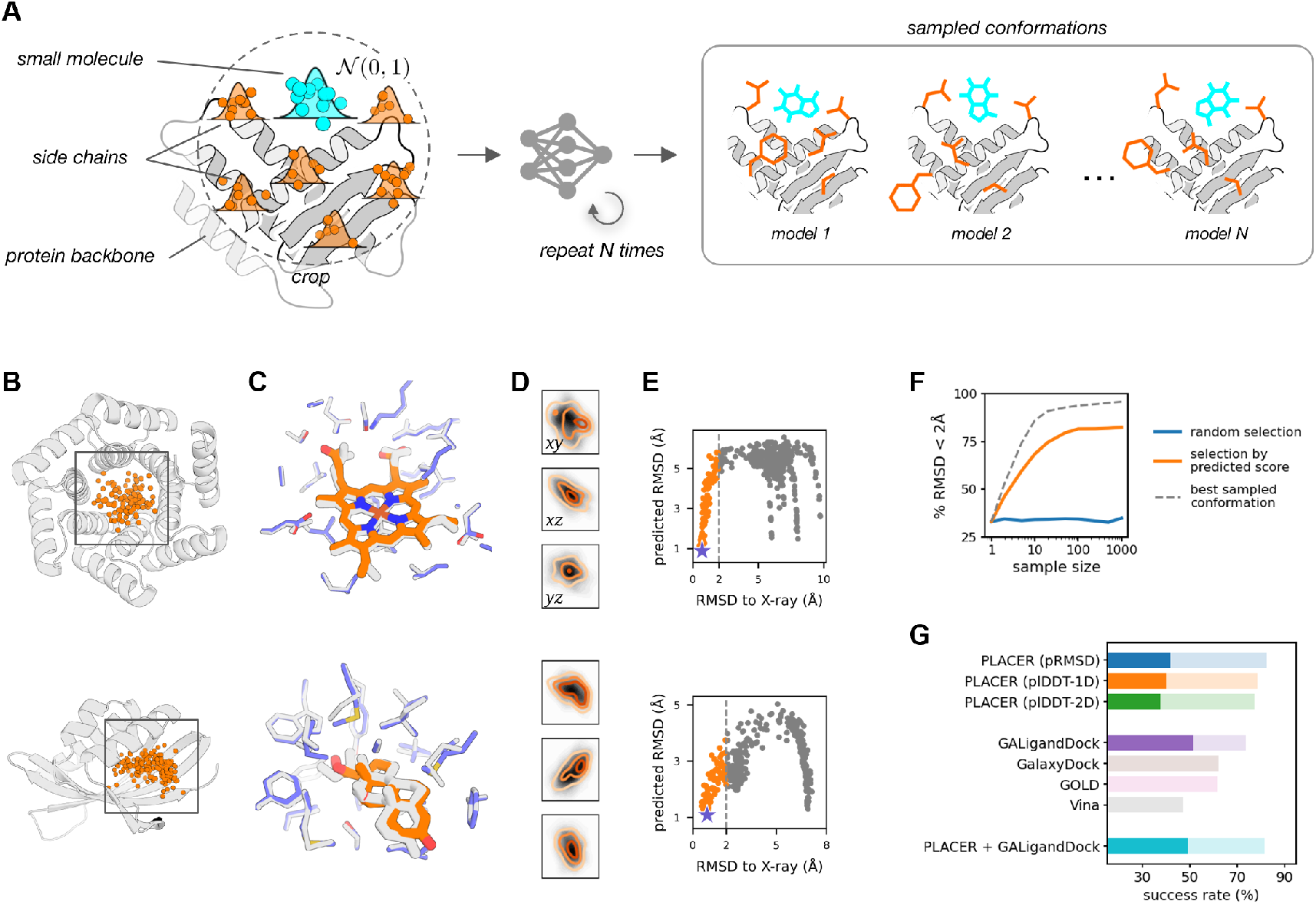
Modeling protein-small molecule interactions with PLACER. **A)** Starting with the protein backbone and the coordinates of the small molecule randomly initialized in the vicinity of the binding site (cyan dots), PLACER predicts protein side chain conformations (which are initially randomized around the respective C-alpha atoms, orange dots) and the structure and the placement of the small molecule relative to the target protein. By repeating this process multiple times PLACER generates a structurally diverse set of docks. **B**,**D)** Sampled starting positions of the small molecule for the two de novo designed small molecule binders. Panel **D** shows projections of the sampled starting points onto the three coordinate planes. Shades of gray show density of all sampled positions; orange contours indicate density of points which resulted in docks with ligand RMSD < 2Å. The two densities largely overlap, suggesting that PLACER is not very sensitive to the initial placement of the ligand in the binding pocket. **C)** Poses with the lowest *pRMSD* scores (blue/orange) closely match the experimental structure (gray). **E)** Models with lower *pRMSD* scores are funneled towards the native conformation demonstrating the discriminative power of the predicted score. **F)** Increasing the sample size and rescoring PLACER models by the *pRMSD* score (orange curve) greatly increases the chance of picking a better model compared to a random selection (blue curve). Scoring by *pRMSD* is however still not optimal (orange vs gray dashed curve). **G)** Among the three accuracy scores predicted by PLACER, *pRMSD* shows best performance (the three bars at the top). Bright and pale bars show docking success rates in the Astex non-native set for ligand RMSD < 1Å and 2Å respectively. The four bars in the middle show performance of the widely used docking tools Vina, GOLD, GalaxyDock, and Rosetta GALigandDock. Sampling docks with PLACER and minimizing and rescoring them using GALigandDock increases docking success rate by 7.3% (ligand RMSD < 1Å) over PLACER alone.

To assess PLACER performance in a more realistic scenario of docking against non-native protein conformations, we used a standard test set of 65 drug targets (22). Each of these targets has a co-crystal structure with a drug molecule, as well as a number of structures either without the drug or with a different small molecule co-crystallized, totaling to 1112 non-native structures over the 65 targets. The docking results highlight the importance of both sampling and scoring aspects of the network (Fig. 3F): the success rate of generating and selecting a near-native conformation of the docked small molecule increases if more models are generated. Among the three confidence prediction metrics tested, the *pRMSD* score shows the best power of selecting near-native conformations (Fig. 3G, top 3 bars), with higher docking success rates than Vina(23), GOLD(22), and GalaxyDock(24) (Fig. 3G, bars in the middle; number were taken from respective references). Compared to the best performing Rosetta GALigandDock method,(25) PLACER performance is higher in the lower accuracy regime (% of complexes with ligand RMSD < 2Å, 82.4% vs 73.6%), but is behind in the high-accuracy regime (RMSD < 1Å, 41.8% vs 51.6%). PLACER performance is still notable however, because, unlike the other methods, the network was not specifically trained on the non-native protein-small molecule docking task. PLACER recreates both the conformations of the small molecule and the protein side chains from scratch, while the other methods tested rely largely on the input protein coordinates. Improved performance can be obtained by combining PLACER with physics-based methods: minimizing the docks generated by the network using the generalized Rosetta force field and estimating binding free energies of the minimized structures gives +7.3% increase in docking success rate at <1Å and no notable change at <2Å; likely due to improved sampling.

### Assessment of enzyme active site design accuracy and pre-organization

A critical challenge in de novo enzyme design is to come up with amino acid sequences that not only encode the target backbone structure but also position the catalytic sidechains and the reaction substrate such that the key sidechain functional groups make the hydrogen bonding and electrostatic interactions with the transition state and each other that are essential to catalysis. As 0.5 Å deviations in distance and 30 degree deviations in angle can have large effects on hydrogen bonding energies, the level of accuracy in positioning required is quite high. Furthermore, the designed active site should be pre-organized in the absence of substrate with the catalytic sidechains largely held in place by intra-protein interactions to reduce the entropic cost of binding and ensure that all catalytic groups are properly positioned; such pre-organization is a notable feature of natural enzyme active sites. The most general way to evaluate such pre-organization is molecular dynamics simulations(26, 27), but as sidechain reconfiguration can occur on the microsecond time scale, such simulations are computationally expensive and cannot be applied to large numbers of designs. Approximate discrete methods such as the Rosetta Rotamer Boltzmann method(28) estimate pre-organization at the individual sidechain level, but are limited by the rotamer approximation and the inability to model coupled and concerted movements of side chains and small molecules. We reasoned that PLACER could enable assessment of the accuracy and pre-organization of designed active sites by generating ensembles of models of small molecules and interacting sidechain placements much more rapidly than molecular dynamics simulations and without the limitations of the Rotamer Boltzmann method.

We applied PLACER to the RA95 series of previously designed retro-aldolase (Fig. 4A) enzymes, and directed evolution generated improved versions of these, for which high resolution crystal structures have been determined(29–31). For every retro-aldolase intermediate shown in Fig. 4A, we ran PLACER 50 times and analyzed the structural diversity of the active site lysine (and covalent adducts of this lysine) through the mean predicted RMSD (*pRMSD*) calculated over side chain atoms and excluding backbone N, Cα, C, O. As illustrated in Figs. 4B, 4C and S5, PLACER generated ensembles are highly varied for the lower activity initial computational designs, indicating a lack of pre-organization, and become increasingly well ordered for the more active evolved versions. This suggests that lack of pre-organization was a major shortcoming of early enzyme design efforts (similar conclusions were reached using considerably more expensive MD simulations(32)), and that PLACER provides a rapid means to assess this, and thus guide enzyme design efforts.

**Figure 4.**
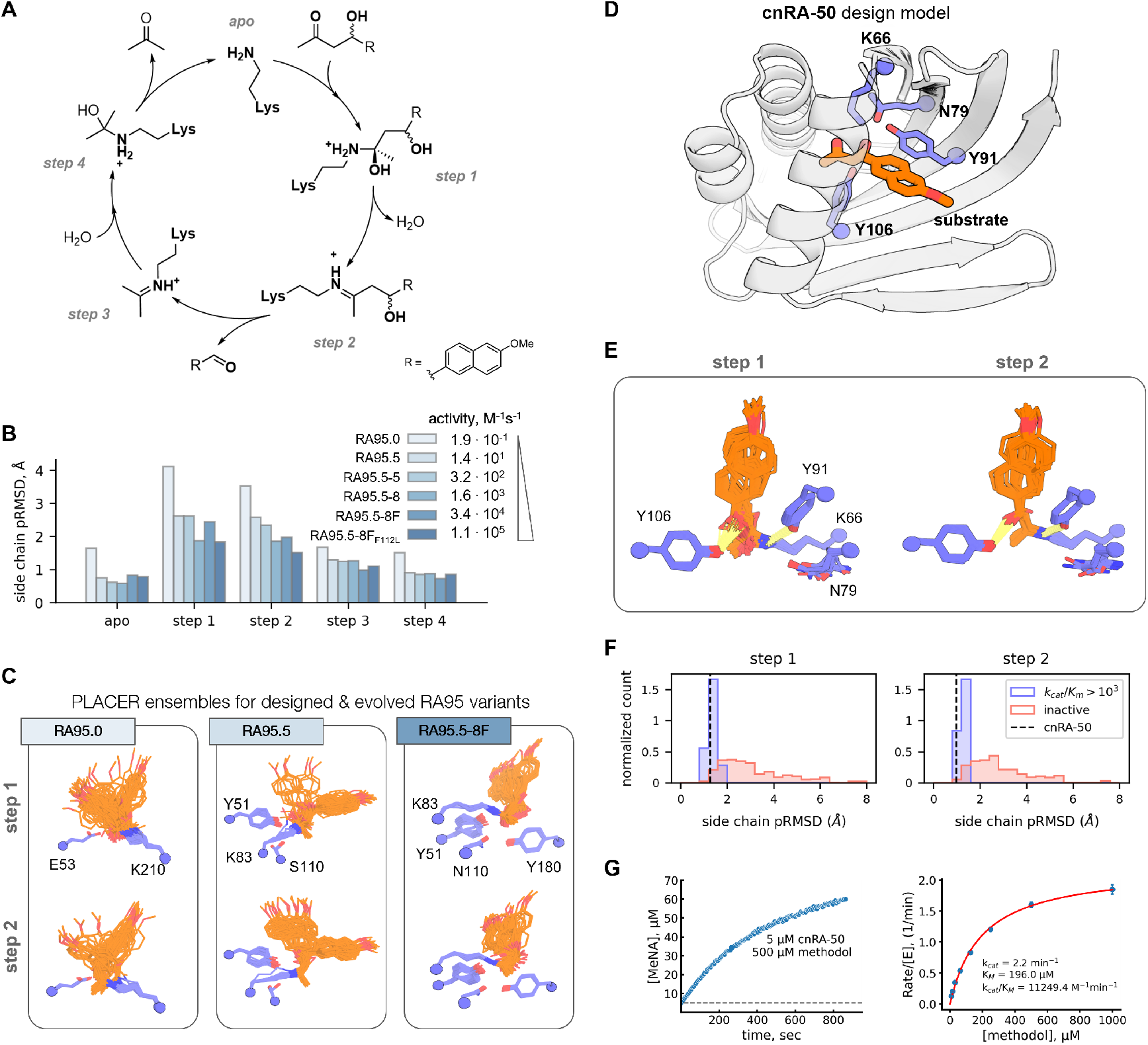
Selecting designs with preorganized catalytic residues increases activity. **A)** Mechanism of the retro-aldol reaction with the key intermediates shown. **B)** Five intermediate states of the RA reaction with methodol were modeled by PLACER for the RA95 series of designs to probe preorganization of the active site lysine and its modifications. **C)** Examples of PLACER ensembles for active sites of the three previously published retro-aldolases with increasing activity show higher degree of preorganization in steps 1 and 2 along the reaction pathway. **D)** Design model of the most active variant, cnRA-50, with the catalytic tetrad highlighted in blue and substrate in orange. **E)** PLACER generated ensembles of lysine-methodol conjugates of catalytic steps 1 and 2 of the active site of cnRA-50. **F)** PLACER active site pre-organization ensemble metrics (pRMSD of conjugated lysine atoms, except backbone) in steps 1 and 2 enrich for selecting more active de novo designed retro-aldolase enzymes. **G)** Retro-aldol reaction catalyzed by cnRA-50: time-course following the formation of 6-methoxy-2-naphthaldehyde (left), and Michaelis-Menten plot (right) obtained from initial velocities.

We further tested the use of PLACER prospectively to guide protein design by making a new round of designed retro-aldolases using deep learning generated NTF2-like folds (21, 33) that previously yielded high activity de novo luciferases(34). We started from an active site description based on the lysine centered catalytic tetrad found in the most active of the evolved retroaldolases(31) (Fig. S3). This active site contains an intricate network of hydrogen bonds keeping the catalytic residues in place (Fig. 4D); we reasoned that PLACER might be helpful to overcome this difficulty by assisting in identification of pre-organized active sites. Designs were generated using RosettaMatch(35) and LigandMPNN(36); full details of the design strategy are provided in the supplemental methods. The activities of the 320 designs were first evaluated in an in vitro transcription translation system (IVTT), enabling rapid identification of promising variants without the need for protein purification. The most active of these were expressed and purified in *E. coli*, and their *k*_cat_/*K*_M_ values determined by measuring activity as a function of substrate concentration. We then examined how PLACER preorganization correlated with activity over all of the tested designs, focusing on the conformational flexibility of the catalytic lysine conjugated to reaction intermediates in the ensembles at each step in the reaction pathway (Fig. 4E). We found that the designs with the highest *k*_cat_/*K*_M_ values were more preorganized than the low activity designs from the initial IVTT screen (Fig. 4F and Fig. S9-11). The most active design, cnRA-50, displayed one of the highest pre-organization levels according to PLACER and had a *k*_cat_/*K*_M_ of 11,000 M^-1^min^-1^ (Fig. 4G), much higher than earlier computational designs prior to directed evolution, and comparable to recent designs made using RFdiffusion and proteinMPNN(37).

## Conclusions

PLACER provides a versatile and rapid approach for generating conformational ensembles of molecules both in isolation and in the context of a protein binding site given the sequence of the protein and the positions of the backbone atoms. Unlike AF3, RoseTTAFold All-Atom, and other protein structure prediction methods, PLACER does not predict protein backbone structure—the benefit is that calculations are considerably more rapid, enabling stochastic generation of conformational ensembles. The use of a consistent atom level representation for all interactions enables facile extension beyond biomolecules to macrocycles and other complex small molecules. The ability to rapidly generate conformational ensembles for protein-small molecule assemblies should be of considerable utility for computational enzyme design and small molecule binder design efforts: the accuracy with which the intended active sites are recapitulated and the extent of pre-organization of the key catalytic/ interacting sidechain functional groups can be readily assessed. PLACER assessment of active site accuracy and pre-organization considerably improves discovery success rates for multi-step serine hydrolases(38) and Zn-dependent metallohydrolases(39), and we describe here the design of retroaldolases with much higher activities than pre-deep learning designs for this reaction, prior to experimental optimization. We anticipate that PLACER based ensemble generation will be broadly useful for modeling the structures of complex non-protein molecules both in isolation and in a protein context, and for evaluating enzyme and protein-small molecule binder designs quite generally.

## Acknowledgments

We thank Luki Goldschmidt and Kandise VanWormer for maintaining the computational and wet lab resources at the Institute for Protein Design. We thank Dr. Declan Evans, and Dr. Florence Hardy for helpful conversations during the development of the method, and Madison A. Kennedy for reading and editing earlier drafts of the manuscript.

## Author Contributions

Designed the research: I.A. and D.B. Developed PLACER architecture and training regimen: I.A. Evaluated PLACER on different prediction tasks: I.A., Y.K., G.Z., S.J.P., A.L., I.K., L.A., G.R.L. Generated and tested retroaldolase designs: Y.K. Contributed code and ideas: I.A., I.K., R.K, J.D. Offered supervision throughout the project: D.B., F.D. Wrote the manuscript: I.A., Y.K, I.K., and D.B. All authors read and contributed to the manuscript.

## Data and Materials Availability

Code and neural network weights are available for download on GitHub: https://github.com/baker-laboratory/PLACER. The model files and sequences of designed retroaldolases are available for download at Zenodo(46). All other data is available in main text or Supplemental Materials.

