## Supplementary Methods for "Modeling protein-small molecule conformational ensembles with PLACER"

**Supplementary Methods** **1**
1. PLACER model training and architecture 2
1.1. Training on the PDB 2
1.2. Training on the CSD 3
1.3. Network input features 3
1.4. PLACER architecture 5
1.5. Losses 5
1.6. List of algorithms 8
1.7. Analysis of potential data leakage for training on the CSD 12
2. Astex non-native benchmark 14
3. Computational analysis of RA95 series of designed and evolved retroaldolases 16
4. Computational de novo design of retroaldolases in NTF2 fold 17
4.1. Identification of focus theozyme configuration 19
4.2. Application of PLACER for design selection 20
4.3. IVTT based retroaldolase activity testing assay 21
4.4. Characterization of individual variants 21
4.5. In depth PLACER analysis of the experimentally tested designs 22
5. References 23

### 1. PLACER model training and architecture

#### 1.1. Training on the PDB

Training and validation sets were compiled from structures deposited to the Protein Data Bank (1) before January 12, 2023. We selected X-ray and EM structures with resolution 2.5Å and better and filtered out entries having ≥10% of unresolved atoms, as well as entries with ≥20 non-standard residues across all polypeptide chains in the asymmetric unit. These filters reduced the size of the set by 33% from 200,713 to 135,172. We additionally removed entries which share similarity to the test set (we used Astex Non-Native Set (2) as our primary small molecule docking benchmark set) either on the protein or the small molecule levels. A PDB structure was discarded if any of its protein chains had ≥30% sequence identity to any of the protein chains in the test set. Structures containing small molecules with ≥80% Tanimoto similarity to the small molecules in the test set were also excluded. We used Open Babel (3) to first compute the small molecule FP4 fingerprints and then to calculate the Tanimoto coefficients. The remaining 119,916 PDB structures were randomly split into training (112,828) and validation (7,090) sets such that none of the protein chains in the two sets share >30% sequence identity. For every structure in the training and validation sets, we counted the number of entries within the set which share ≥30% sequence identity on the protein chain level. The inverse of these counts were then used during training and validation to sample the structures from the respective set.

During training and validation, we do not split the PDB entries into individual chains or molecules but operate on the whole asymmetric units. We directly parse PDBx/mmCIF files keeping all molecular content; only water molecules are excluded. The Chemical Component Dictionary (4) is used to construct chemical graphs for individual residues and small molecules observed in a PDB entry; nodes in a chemical graph represent individual atoms (only non-hydrogen atoms are considered) and edges represent chemical bonds between atoms. The residue-level graphs are joined together into a single chemical graph for the whole PDB entry. Edges are added to connect subgraphs representing individual residues within a polymer chain (proteins, DNA, RNA), as well as are edges parsed from the `struct_conn` data records of the PDBx/mmCIF file. Atomic coordinates are parsed from the PDBx/mmCIF file and are saved on a corresponding node of the chemical graph.

PDB structures were cropped to at most 600 heavy atoms to fit the GPU memory. First, we split all atoms in the input structure into four classes (protein, nucleic acid, small molecule, metal), then randomly choose one of these classes with ratios 1:1:5:1, and finally sample a random heavy atom from the selected class to be the center of the cropped region. The crop is defined as the collection of residues and small molecules in the PDB structure which are closest in space to the residue or the small molecule the crop center belongs to. Atom coordinates in the crop undergo corruption. First, we collect all atoms in the 8-hop neighborhood of the crop center in the chemical graph and add them to the corruption set. Backbone atoms in polypeptide (N,CA,C), polyribo- and polydeoxyribonucleotide chains (O3', C3', C4', C5', O5', P) which are not part of the corruption set are kept fixed with only a minor Gaussian noise ($\sigma$ = 0.1Å) added to the atom coordinates to make the network more robust to small displacements in atomic coordinates of the backbone atoms(5). All the remaining non-backbone atoms are included in the corruption set. The chemical graph corresponding to the crop is then split into connected components. If a connected component has fixed backbone atoms, then the corrupted atoms in that component are initialized with the coordinates of the closest backbone atom in the graph (based on the shortest path in the graph). Otherwise, a random atom is picked from the component, Gaussian noise with $\sigma$ = 1.5Å is added to its coordinates, and all other atoms reachable from the selected atom are collapsed to it. Finally, Gaussian noise with $\sigma$ = 1.5Å is applied to all atoms in the corruption set.

#### 1.2. Training on the CSD

The PDB-derived datasets were complemented by the crystal structures of small molecules deposited to the Cambridge Structural Database (CSD v5.43; November 2021)(6). We used structures with R-factor 7.5% and better and no disorder and excluded polymeric and organo-metallic entries; the number of heavy atoms per small molecule was limited to the 3..80 range. Only molecules which could be successfully parsed by Open Babel were retained. CSD structures with multiple molecules in the asymmetric unit were split into individual molecules, followed by merging of identical molecules with the same canonical SMILES string into one entry for training and validation. Upon merging, we kept 3D coordinates of all the instances to account for alternative conformations during training. Training and validation splits were done using 75% Tanimoto similarity cutoff, yielding 226,684 and 7,116 examples, respectively. During training, molecules were sampled with frequencies inversely proportional to the number of similar molecules in the set at Tanimoto similarity 75%.

#### 1.3. Network input features

Training examples from both the PDB and the CSD were featurized in the same way. The information about atom and bond types which were used as inputs to PLACER is summarized in Table S1. In addition to atom single (`f1d`) and pair (`f2d`) features, PLACER also utilizes information about the local topology of the input molecules which is represented by bonding, bond length, chirality and planarity terms (for atoms which are immediately connected in the chemical graph, Table S1, entries 18-21).

**Table S1**. Description of PLACER input features.

| Entry | Input name (Shape) | Description |
|---|---|---|
| 1 | **`xyz`** ($N_{atoms}$, 3) | initial atomic coordinates |
| 2 | **`group`** ($N_{atoms}$, 7) | element's group in the periodic table, 1-hot encoded |
| 3 | **`period`** ($N_{atoms}$, 18) | element's period in the periodic table, 1-hot encoded |
| 4 | **`is_lanthanide`** ($N_{atoms}$, 1) | a boolean flag indicating whether the element is a lanthanide |
| 5 | **`is_actinide`** ($N_{atoms}$, 1) | a boolean flag indicating whether the element is an actinide |
| 6 | **`lanthanide_group`** ($N_{atoms}$, 15) | element's group within lanthanides, 1-hot encoded |
| 7 | **`actinide_group`** ($N_{atoms}$, 15) | element's group within actinides, 1-hot encoded |

| | | |
|---|---|---|
| **8** | **`charge`**<br>($N_{atoms}$, 7) | total charge of the atom, 1-hot encoded from set {-3,-2,-1,0,1,2,3} |
| **9** | **`nhyd`**<br>($N_{atoms}$, 4) | number of hydrogens connected to the atom, 1-hot encoded |
| **10** | **`hyb`**<br>($N_{atoms}$, 6) | hybridization state of the atom, 1-hot encoded from set {*sp*, $sp^2$, $sp^3$, square planar, trigonal bipyramidal, octahedral} |
| **11** | **`is_corrupted`**<br>($N_{atoms}$, 1) | a boolean flag indicating whether the atom belongs to the corruption set |
| **12** | **`f1d`**<br>($N_{atoms}$, 75) | concatenation of **`group`**, **`period`**, **`is_lanthanide`**, **`is_actinide`**, **`lanthanide_group`**, **`actinide_group`**, **`charge`**, **`nhyd`**, **`hyb`**, **`is_corrupted`** |
| **13** | **`is_aromatic`**<br>($N_{atoms}$, $N_{atoms}$, 2) | a boolean flag indicating whether the bond is aromatic |
| **14** | **`is_in_ring`**<br>($N_{atoms}$, $N_{atoms}$, 2) | a boolean flag indicating whether the bond is in a ring |
| **15** | **`bond_order`**<br>($N_{atoms}$, $N_{atoms}$, 4) | bond order, 1-hot encoded from set {single, double, triple, other} |
| **16** | **`bond_separation`**<br>($N_{atoms}$, $N_{atoms}$, 7) | an integer counting the number of covalent bonds between any two atoms in the chemical graph, 1-hot encoded. If number of bonds between atoms >7, then separation=0. |
| **17** | **`f2d`**<br>($N_{atoms}$, $N_{atoms}$, 15) | concatenation of **`is_aromatic`**, **`is_in_ring`**, **`bond_order`**, **`bond_separation`** |
| **18** | $\mathbf{S}_{\mathbf{bonds}}$<br>($N_{bonds}$, 2) | pairs of atoms connected by a chemical bond |
| **19** | **`bond_len`**<br>($N_{bonds}$) | equilibrium bond lengths for bonds in $\mathbf{S}_{\mathbf{bonds}}$ based on the covalent radii and bond order (from Open Babel) |
| **20** | $\mathbf{S}_{\mathbf{chirals}}$<br>($N_{chirals}$, 4) | ($o$,$i$,$j$,$k$) tuples where $o$ is the index of the central pyramidal or tetrahedral $sp^3$ hybridized atom and its neighboring atoms $i$,$j$,$k$ ordered clockwise |
| **21** | $\mathbf{S}_{\mathbf{planars}}$<br>($N_{planars}$, 4) | ($o$,$i$,$j$,$k$) tuples for planar geometries around an $sp^2$ hybridized atom $o$ and its neighboring atoms $i$,$j$,$k$ |

#### 1.4. PLACER architecture

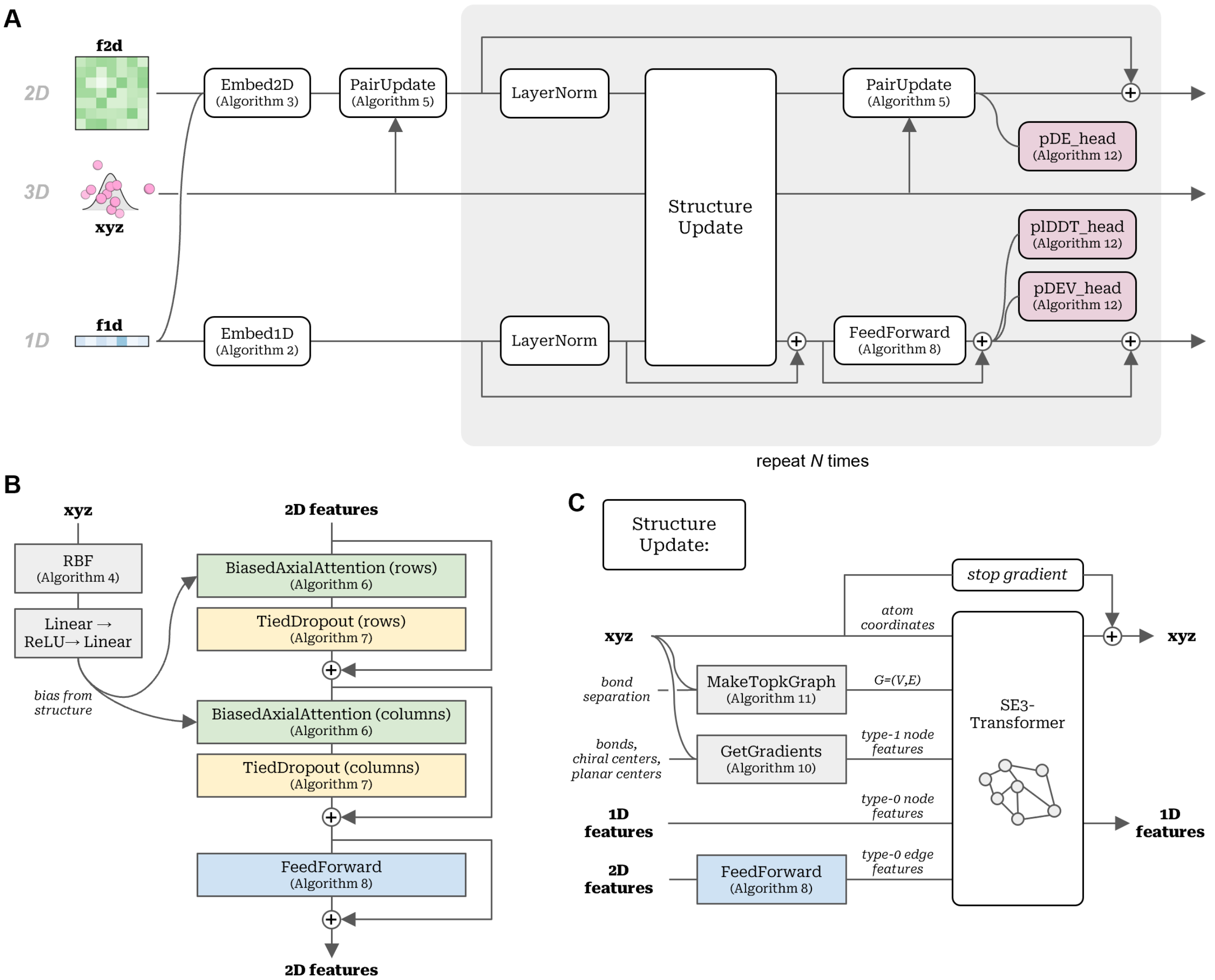


**Figure S1**. **PLACER architecture**. **A)** The three-track network schematics (Algorithm 1). **B)** PairUpdate module (Algorithm 5). **C)** Structure update module.

#### 1.5. Losses

We use a combination of structure losses and confidence prediction losses to train PLACER:

$$L = w_{aaFAPE}L_{aaFAPE} + w_{aaFAPE}^{interaction}L_{aaFAPE}^{interaction} + w_{aaFAPE}^{small\ molecule}L_{aaFAPE}^{small\ molecule} + \quad (1)$$

$$w_{bond}L_{bond} + w_{angle}L_{angle} + w_{chiral}L_{chiral} + w_{planar}L_{planar} +$$

$$w_{plDDT}L_{plDDT} + w_{pDE}L_{pDE} + w_{pRMSD}L_{pRMSD}$$

**Table S2**. Loss weights used during PLACER training.

| training stage | $w_{aaFAPE}$ | $w_{aaFAPE}^{interaction}$ | $w_{aaFAPE}^{small\ molecule}$ | $w_{bond}$ | $w_{angle}$ | $w_{chiral}$ | $w_{planar}$ | $w_{plDDT}$ | $w_{pDE}$ | $w_{pRMSD}$ |
|---|---|---|---|---|---|---|---|---|---|---|
| 1 | 1.0 | 0.0 | 1.0 | 0.0 | 0.0 | 0.0 | 0.0 | 1.0 | 1.0 | 1.0 |
| 2 | 1.0 | 1.0 | 1.0 | 1.0 | 1.0 | 1.0 | 1.0 | 1.0 | 1.0 | 1.0 |

**All-atom FAPE loss** $L_{aaFAPE}$ is a modification of the original FAPE loss introduced in Ref. (7) allowing to operate on molecules of any type and to treat the entire molecule flexibly (no need to define rigid groups, like backbone N-Ca-C in the original FAPE). First, we enumerate all triples $i, j, k$ of atoms which are covalently bonded (there are covalent bonds between atoms $i, j$ and $j, k$). Spatial coordinates $\vec{x}_i$,$\vec{x}_j$,$\vec{x}_k$ of these three atoms in the predicted structure are used to construct a local reference frame according to Eqs. 2:

$$\vec{u} = \vec{x}_j - \vec{x}_i \quad (2)$$

$$\vec{v} = \vec{x}_k - \vec{x}_j$$

$$\vec{e}_1 = \frac{\vec{u}\times\vec{v}}{|\vec{u}\times\vec{v}|}, \vec{e}_3 = \frac{\vec{u}-\vec{v}}{|\vec{u}-\vec{v}|}, \vec{e}_2 = \vec{e}_3 \times \vec{e}_1$$

$$R^x_{(ijk)} = [\vec{x}_i, \vec{x}_j, \vec{x}_k]$$

Similarly, a frame $R^y_{(ijk)}$ is constructed from the atom coordinates in the reference experimental structure. The two frames $R^x_{(ijk)}$, $R^y_{(ijk)}$ and the two sets of atom coordinates – one in the model structure $\{\vec{x}\}$ and one in the reference $\{\vec{y}\}$ – are then used to calculate deviations in atom positions once the frames are aligned:

$d_{(ijk),l} = \left|R^x_{(ijk)}(\vec{x}_l - \vec{x}_j) - R^y_{(ijk)}(\vec{y}_l - \vec{y}_j)\right|$ – deviation in positions for atom $l$

$$L_{aaFAPE} = \frac{1}{N_{atoms}N_{frames}} \sum_{(ijk)\in S_{frames}} \sum_{l=1}^{N_{atoms}} d_{(ijk),l}$$

To put more weight during training on the small molecule, and the interaction between the small molecule and the rest of the structure, we also used two slightly modified versions of all-atom FAPE with ${L_{aaFAPE}}^{p,sm}$ measuring deviations of small molecule atoms in the protein frames and ${L_{aaFAPE}}^{sm,sm}$ emphasizing the internal structure of the small molecule.

$$ {L_{aaFAPE}}^{interaction} = L_{aaFAPE}(S_{frames} \in protein, l \in small\ molecule)$$
$$ {L_{aaFAPE}}^{small\ molecule} = (S_{frames} \in small\ molecule, l \in small\ molecule)$$

**Bonded losses.** Bond loss term $L_{bond}$ measures the distance deviation for covalently bonded atoms between the model and the reference structures:

$$L_{bond} = MSE_{(ij)\in S_{bonds}}(|\vec{x_i} - \vec{x_j}|, |\vec{y_i} - \vec{y_j}|)$$

where $MSE$ is a mean squared error loss. $MSE$ loss is also used to measure deviations in bonded angles:

$$L_{angle} = MSE_{(ijk)\in S_{angles}}\left(\alpha^x_{(ijk)}, \alpha^y_{(ijk)}\right)$$

where $\alpha^x_{(ijk)}$ and $\alpha^y_{(ijk)}$ are planar angles (in radians) between atoms $i, j, k$ calculated from the model and the reference structures respectively.

Additional loss term $L_{chiral}$ is imposed on all quadruples of atoms $o, i, j, k \in S_{chirals}$ where the three atoms $i, j, k$ are covalently bonded to an $sp^3$-hybridized central atom $o$ to force the off-plane geometry. This is achieved by first computing the angle $\theta_{(oijk)}$ between the normal to the $o, i, j$ plane $\vec{n}_{(oij)} = (\overrightarrow{x_i} - \overrightarrow{x_o}) \times (\overrightarrow{x_j} - \overrightarrow{x_o})$ and the off-plane vector $\vec{v}_{(ok)} = \overrightarrow{x_k} - \overrightarrow{x_o}$ and then applying the $MSE$ loss:

$$L_{chiral} = MSE_{(oijk)\in S_{chirals}}\left(\theta^x_{(oijk)}, \theta^y_{(oijk)}\right)$$

A similar loss term $L_{planar}$ is used to enforce planar geometry around the $sp^2$-hybridized atom $o$ and its neighboring atoms $i, j, k$:

$$L_{planar} = MSE_{(oijk)\in S_{planars}}\left(\theta^x_{(oijk)}, \theta^y_{(oijk)}\right)$$

**Confidence losses.** We model per-atom confidences in two different ways. One is through predicted lDDT (plDDT) score which has previously been introduced in the context of the AF2 network(7); for PLACER we extended it to the all atom representation. Additionally, we predict uncertainties in atomic positions through estimating the expected standard deviations between atom $x, y, z$ coordinates in the model and in the reference structures. These standard deviations, if combined over the entire structure or a region of interest, give rise to the predicted RMSD (pRMSD) score (see Main text for details). Lastly, to estimate atom-atom pair accuracies, we predicted errors in distances between every two atoms. All-by-all distance matrix of the model was subtracted from the one of the reference to get the signed distance error matrix (referred to as "estograms").(8) This matrix was then binned into 102 bins with 100 equal bins covering the [-5;5] interval plus the two bins corresponding to the outside regions (-∞;-5) and (5;∞) ). The network was trained to make categorical predictions over these bins for which the categorical cross-entropy loss was used during training.

#### 1.6. List of algorithms

**Algorithm 1.** PLACER network

```
def PLACER(X_i, f1d_i, f2d_i,j, separation, bonds, bondlen, chirals, planars, N_iter):
# embed inputs
    single_i^prev = Embed1D(f1d_i, c = 64)                                  # Algorithm 2, single_i ∈ ℝ^64
    pair_i,j^prev = Embed2D(f2d_i,j, c = 128)                               # Algorithm 3,  pair_i,j ∈ ℝ^128
    pair_i,j^prev = PairUpdate(pair_i,j^prev, X_i, c = 32, N_head = 4, p_drop = 0.15)   # Algorithm 5
# iteratively update coordinates and 1D and 2D embeddings
    Xs, plDDTs, pDEVs, pDEs = [X_i], [ ], [ ], [ ]                          # lists for saving results
    scale_factor = 100.0
    for iter = 1 ... N_iter:                                    # network weights are shared within this block
        single_i = LayerNorm(single_i^prev) if iter > 1 else single_i^prev
        pair_i,j = LayerNorm(pair_i,j^prev) if iter > 1 else pair_i,j^prev
        g_i = GetGradients(Xs[iter], bonds, bondlen, chirals, planars)  # Algorithm 10,  g_i ∈ ℝ^3,3
        G = MakeTopkGraph(Xs[iter], separation, topk = 32)              # Algorithm
11
        edge_features_k = FeedForward(pair_i,j ∈ G.edges, normalize=False)   # Algorithm 8
        state_i , dx_i = SE3_Transformer(G, single_i, l1_i, edges_k)
# update coords and embeddings
        X_i = stopgrad(X_i) + dx_i / scale_factor
        single_i = UpdateSingle(single_i + state_i)     # FeedForward(x,c=128,n=2,pdrop=0.15,normalize=True)
        pair_i,j = PairUpdate(pair_i,j, X_i, c = 32, N_head = 4, p_drop = 0.15)
# auxiliary heads
        plddt_i = PredictError(single_i, bins=51)                        # Algorithm 12
        dev_i = PredictError(single_i, bins=1)                           # Algorithm 12
        pde_i,j = PredictError(pair_i,j+pair_j,i, bins=102)              # Algorithm 12
# residual connections
        single_i^prev += single_i
        pair_i,j^prev += pair_i,j
# append predictions from the current iteration to the output lists
        Xs += [X_i]
        plDDTs += [plddt_i]
        pDEVs += [dev_i]
        pDEs += [pde_i,j]
    return Xs, pDEs, plDDTs, pDEVs
```

**Algorithm 2.** Initial embedding of node features

```
def Embed1D(f1d_i, c = 64):
    s_i = Linear(f1d_i)                          # s_i ∈ ℝ^c
    s_i = LayerNorm(Linear(ReLU(s_i)))
    return s_i
```

**Algorithm 3.** Initial embedding of pair features

**def** Embed2D($\mathbf{f1d}_i$, $\mathbf{f2d}_{i,j}$, $c$ = 128):

$\mathbf{s}_i^{left}$, $\mathbf{s}_i^{right}$ = Linear($\mathbf{f1d}_i$) # $\mathbf{s}_i^{left}, \mathbf{s}_i^{right} \in \mathbb{R}^c$
$\mathbf{p}_{i,j}$ = Linear($\mathbf{f2d}_{i,j}$) # $\mathbf{p}_{i,j} \in \mathbb{R}^c$
$\mathbf{s}_{i,j} = \mathbf{s}_i^{left} + \mathbf{s}_j^{right}$ # *outer summation*
$\mathbf{p}_{i,j}$ = concat($\mathbf{p}_{i,j}$, $\mathbf{s}_{i,j}$) # $\mathbf{p}_{i,j} \in \mathbb{R}^{2c}$
$\mathbf{p}_{i,j}$ = ReLU($\mathbf{p}_{i,j}$)
$\mathbf{p}_{i,j}$ = Linear($\mathbf{p}_{i,j}$) # $\mathbf{p}_{i,j} \in \mathbb{R}^c$
$\mathbf{p}_{i,j}$ = LayerNorm($\mathbf{p}_{i,j}$)
**return** $\mathbf{p}_{i,j}$

**Algorithm 4.** Project pairwise distances for a set of points onto Gaussian radial basis functions

**def** RBF($\mathbf{X}_i$, $N_{rbf}$ = 32, $D_{max}$ = 20.0):

$D_{i,j} = \|\mathbf{X}_i - \mathbf{X}_j\|$
$\sigma = D_{max} / N_{rbf}$
$\mathbf{rbf}_{i,j}$ = concat(exp(−(($D_{i,j} - k\sigma)/\sigma))^2$) **for** $k$ **in** 1,...,$N_{rbf}$) # $\mathbf{rbf}_{i,j} \in \mathbb{R}^{Nrbf}$
**return** $\mathbf{rbf}_{i,j}$

**Algorithm 5.** Pair update with bias from structure

**def** PairUpdate($\mathbf{pair}_{i,j}$, $\mathbf{xyz}_i$, $c$ = 32, $N_{head}$ = 4, $p_{drop}$ = 0.15):

$\mathbf{rbf}_{i,j}$ = RBF($\mathbf{xyz}_i$, $N_{rbf} = c$) # Algorithm 4, $\mathbf{rfb}_{i,j} \in \mathbb{R}^c$
$\mathbf{bias}_{i,j}$ = Linear($\mathbf{rbf}_{i,j}$) # $\mathbf{bias}_{i,j} \in \mathbb{R}^c$
$\mathbf{bias}_{i,j}$ = Linear(ReLU($\mathbf{bias}_{i,j}$)) # $\mathbf{bias}_{i,j} \in \mathbb{R}^{cn}$
*# attention and dropout applied to rows*
$\mathbf{pair}_{i,j}^{row}$ = BiasedAxialAttention($\mathbf{pair}_{i,j}$, $\mathbf{bias}_{i,j}$, $c$, $N_{head}$, is_row = *True*) # $\mathbf{pair}_{i,j}^{row} \in \mathbb{R}^{cn}$
$\mathbf{pair}_{i,j}^{row}$ = TiedDropout($\mathbf{pair}_{i,j}^{row}$, $p_{drop}$, is_row = *True*)
$\mathbf{pair}_{i,j}$ += $\mathbf{pair}_{i,j}^{row}$
*# attention and dropout applied to columns*
$\mathbf{pair}_{i,j}^{col}$ = BiasedAxialAttention($\mathbf{pair}_{i,j}$, $\mathbf{bias}_{i,j}$, $c$, $N_{head}$, is_row = *False*) # $\mathbf{pair}_{i,j}^{col} \in \mathbb{R}^{cn}$
$\mathbf{pair}_{i,j}^{col}$ = TiedDropout($\mathbf{pair}_{i,j}^{col}$, $p_{drop}$, is_row = *False*)
$\mathbf{pair}_{i,j}$ += $\mathbf{pair}_{i,j}^{col}$
*# feed-forward layer*
$\mathbf{pair}_{i,j}$ += FeedForward($\mathbf{pair}_{i,j}$, $p_{drop}$) # $\mathbf{pair}_{i,j} \in \mathbb{R}^{cn}$
**return** $\mathbf{pair}_{i,j}$

**Algorithm 6.** Tied axial attention with bias from structure

**def** BiasedAxialAttention($\mathbf{pair}_{i,j}$, $\mathbf{bias}_{i,j}$, $c$, $N_{head}$, is_row = *True*):
 **if** is_row **is** *True*: # *flip rows and columns*
  $\mathbf{pair}_{i,j} = \mathbf{pair}_{j,i}$
  $\mathbf{bias}_{i,j} = \mathbf{bias}_{j,i}$
 $\mathbf{pair}_{i,j}$ = LayerNorm($\mathbf{pair}_{i,j}$)
 $bias_{i,j}^h$ = Linear(LayerNorm($\mathbf{bias}_{i,j}$)) # $bias_{i,j}^h \in \mathbb{R}$, $h = 1 \ldots N_{head}$
 $\mathbf{gate}_{i,j}^h$ = sigmoid(Linear($\mathbf{pair}_{i,j}$)) # $\mathbf{gate}_{i,j}^h \in \mathbb{R}^c$
 $\mathbf{q}_{i,j}^h, \mathbf{k}_{i,j}^h, \mathbf{v}_{i,j}^h$ = Linear($\mathbf{pair}_{i,j}$, bias = *False*) # $\mathbf{q}_{i,j}^h, \mathbf{k}_{i,j}^h, \mathbf{v}_{i,j}^h \in \mathbb{R}^c$
 $\mathbf{q}_{i,j}^h = \mathbf{q}_{i,j}^h / c^{½}$
 $\mathbf{k}_{i,j}^h = \mathbf{k}_{i,j}^h / N_{atoms}^{½}$
 $attn_{i,j}^h = \text{softmax}_j(\text{sum}_n((\mathbf{q}_{n,i}^h)^{\top} \mathbf{k}_{n,j}^h) + bias_{i,j}^h)$ # *tied attention with bias*; $attn_{i,j}^h \in \mathbb{R}$
 $\mathbf{out}_{i,j}^h = \text{sum}_k(attn_{i,k}^h \, \mathbf{v}_{j,k}^h)$
 $\mathbf{out}_{i,j} = \text{flatten}_h(\mathbf{out}_{i,j}^h \odot \mathbf{gate}_{i,j}^h)$
 **if** is_row **is** *True*: $\mathbf{out}_{i,j} = \mathbf{out}_{j,i}$ # *flip rows and columns back*
 **return** $\mathbf{out}_{i,j}$

**Algorithm 7.** Dropout layer to zero out entire rows or columns from the pair representation

**def** TiedDropout($\mathbf{pair}_{i,j}$, $p_{drop}$ = 0.15, is_row = *True*):
 $mask_i \sim$ Bernoulli($1 - p_{drop}$) # *sample from Bernoulli distribution*
 **if** is_row **is** *True*:
  $\mathbf{pair}_{i,j} = mask_i \cdot \mathbf{pair}_{i,j} / (1 - p_{drop})$ # *drop row*
 **else**:
  $\mathbf{pair}_{i,j} = mask_j \cdot \mathbf{pair}_{i,j} / (1 - p_{drop})$ # *drop column*
 **return** $\mathbf{pair}_{i,j}$

**Algorithm 8.** Feed-forward layer

**def** FeedForward($\mathbf{x}$, $c$ = 128, $n$ = 2, $p_{drop}$ = 0.10, normalize = *True*):
 **if** normalize **is** *True*: $\mathbf{x}$ = LayerNorm($\mathbf{x}$)
 $\mathbf{x}$ = Linear($\mathbf{x}$) # $\mathbf{x} \in \mathbb{R}^{cn}$ *(project up)*
 $\mathbf{x}$ = Linear(Dropout(ReLU($\mathbf{x}$, $p_{drop}$))) # $\mathbf{x} \in \mathbb{R}^c$ *(project down)*
 **return** $\mathbf{x}$

**Algorithm 9.** Triple product of three vectors

**def** TripleProduct($\mathbf{o}$, $\mathbf{a}$, $\mathbf{b}$, $\mathbf{c}$): # $\mathbf{o},\mathbf{a},\mathbf{b},\mathbf{c} \in \mathbb{R}^3$
### *input vectors are defined by 4 points;* **o** *is the* initial point of the three vectors, **a**,**b**,**c** are endpoints
 $v = (\mathbf{o} - \mathbf{a}) \cdot ((\mathbf{o} - \mathbf{b}) \times (\mathbf{o} - \mathbf{c}))$ # $v \in \mathbb{R}$

**return** $v$

---

**Algorithm 10.** Get gradients of bonded geometry w.r.t. atom coordinates

**def** GetGradients($\mathbf{X}_i$, bonds, bondlen, chirals, planars):

$\mathbf{X}_i$ = stopgrad($\mathbf{X}_i$)

$L^{bond} = \text{mean}_{i,j\,\in\,\text{bonds}}\,(\lVert\mathbf{X}_i - \mathbf{X}_j\rVert - \text{bondlen}_{(i,j)})^2$ # $L^{bond}, L^{chiral}, L^{planar} \in \mathbb{R}$

$L^{chiral} = \text{mean}_{o,i,j,k\,\in\,\text{chirals}}\,(\text{TripleProduct}(\mathbf{X}_o,\mathbf{X}_i,\mathbf{X}_j,\mathbf{X}_k) - 0.70710678)^2$

$L^{planar} = \text{mean}_{o,i,j,k\,\in\,\text{planars}}\,(\text{TripleProduct}(\mathbf{X}_o,\mathbf{X}_i,\mathbf{X}_j,\mathbf{X}_k))^2$

*# we use pytorch automatic differentiation to calculate partial derivatives*

$\mathbf{g}_i^{bond} = \partial L^{bond}/\partial\mathbf{X}_i$ # $\mathbf{g}_i^{bond}, \mathbf{g}_i^{chiral}, \mathbf{g}_i^{planar} \in \mathbb{R}^3$

$\mathbf{g}_i^{chiral} = \partial L^{chiral}/\partial\mathbf{X}_i$

$\mathbf{g}_i^{planar} = \partial L^{planar}/\partial\mathbf{X}_i$

$\mathbf{g}_i$ = concat($\mathbf{g}_i^{bond}, \mathbf{g}_i^{chiral}, \mathbf{g}_i^{planar}$) # $\mathbf{g}_i \in \mathbb{R}^{3\times 3}$

return $\mathbf{g}_i$

---

**Algorithm 11.** Create *topk* neighbor graph

**def** MakeTopkGraph($\mathbf{X}_i$, separation, topk = 32):

edges = ∅

$D_{i,j} = \lVert\mathbf{X}_i - \mathbf{X}_j\rVert$

*# for every node i, collect bonded and non-bonded neighbors*

**for all** $i = 1,\ldots,N_{atoms}$:

  $n_i^{bonded}$ = {topk/2 $j$'s with smallest separation$_{i,j}$ and $i \neq j$}

  $n_i^{non\text{-}bonded}$ = {topk/2 $j$'s with smallest $D_{i,j}$ which are not in $n_i^{bonded}$, $i \neq j$}

  $n_i = n_i^{bonded} \cup n_i^{non\text{-}bonded}$

  edges = edges ∪ {$(i,j)$ **for** $j$ **in** $n_i$}

*# create a graph**

$\boldsymbol{G} \leftarrow$ graph object with $V = 1,\ldots,N_{atoms}$ and $E$ = edges

**return** $\boldsymbol{G}$

---

* Internally, we use DGL library to represent the graph in the SE3-Transformer module

---

**Algorithm 12.** Error prediction heads (plDDT, pDE, pDEV)

**def** PredictError($\mathbf{x}$, c=256, p_drop=0.0, bins=51 ):

$\mathbf{x}$ = LayerNorm($\mathbf{x}$)

$\mathbf{x}$ = Linear($\mathbf{x}$) # $\mathbf{x} \in \mathbb{R}^c$

$\mathbf{x}$ = Linear(Dropout(ReLU($\mathbf{x}$, $p_{drop}$))) # $\mathbf{x} \in \mathbb{R}^c$

$\boldsymbol{\varepsilon}$ = Linear(Dropout(ReLU($\mathbf{x}$, $p_{drop}$))) # $\boldsymbol{\varepsilon} \in \mathbb{R}^{bins}$

**return** $\boldsymbol{\varepsilon}$

**Algorithm 13.** All-atom FAPE loss

**def** GetFrame($\mathbf{a}$, $\mathbf{b}$, $\mathbf{c}$):
$\mathbf{u} = \mathbf{b} - \mathbf{a}$
$\mathbf{v} = \mathbf{c} - \mathbf{b}$
$\mathbf{e}_3 = (\mathbf{u} - \mathbf{v}) / \|\mathbf{u} - \mathbf{v}\|$
$\mathbf{e}_1 = \mathbf{v} \times \mathbf{u} / \|\mathbf{v} \times \mathbf{u}\|$
$\mathbf{e}_2 = \mathbf{e}_3 \times \mathbf{e}_1$
$\mathbf{R}$ = concat($\mathbf{e}_1$ , $\mathbf{e}_2$, $\mathbf{e}_3$)
**return R**

**Algorithm 14.** All-atom FAPE loss

**def** FAPE($\{\mathbf{X}_i\}$, $\{\mathbf{Y}_i\}$, $S_{frames}$, $\delta$=1.0, cut=10.0):
**for all** $i = 1,\ldots,N_{atoms}$:
**for** $k,l,m$ **in** $S_{frames}$:
$\mathbf{R}^{\mathbf{X}} = \mathbf{GetFrame}(\mathbf{X}_k, \mathbf{X}_l, \mathbf{X}_m)$
$\mathbf{R}^{\mathbf{Y}} = \mathbf{GetFrame}(\mathbf{Y}_k, \mathbf{Y}_l, \mathbf{Y}_m)$
$\{\mathbf{X}_i\}^{\mathrm{rot}} = \mathbf{R}^{\mathbf{X}} (\{\mathbf{X}_i\} - \mathbf{X}_l)$
$\{\mathbf{Y}_i\}^{\mathrm{rot}} = \mathbf{R}^{\mathbf{Y}} (\{\mathbf{Y}_i\} - \mathbf{Y}_l)$
$\{d_i\} = \{\|\mathbf{X}_i - \mathbf{Y}_i\|\}$
$v_i = \frac{1}{2}\{d_i\}^2$ if $|\{d_i\}| < \delta$
$v_i$ += $\delta * (\{d_i\} - \frac{1}{2}\delta)$ if $|\{d_i\}| >= \delta$
$v_i = clamp(v_i, cut)$
**return** $\mathrm{mean}_i(v_i)$

#### 1.7. Analysis of potential data leakage for training on the CSD

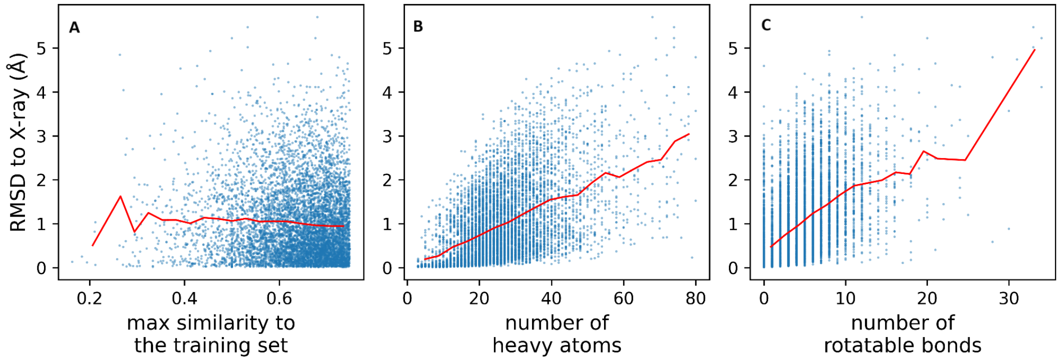


**Figure S2**. **Analysis of PLACER performance on the CSD test set**. **A)** Performance on the 7,116 test set examples is almost independent of the highest Tanimoto similarity to the test set. Factors like molecule size **(B)** and flexibility **(C)** on the other hand negatively affect PLACER performance. Trend lines are shown in red.

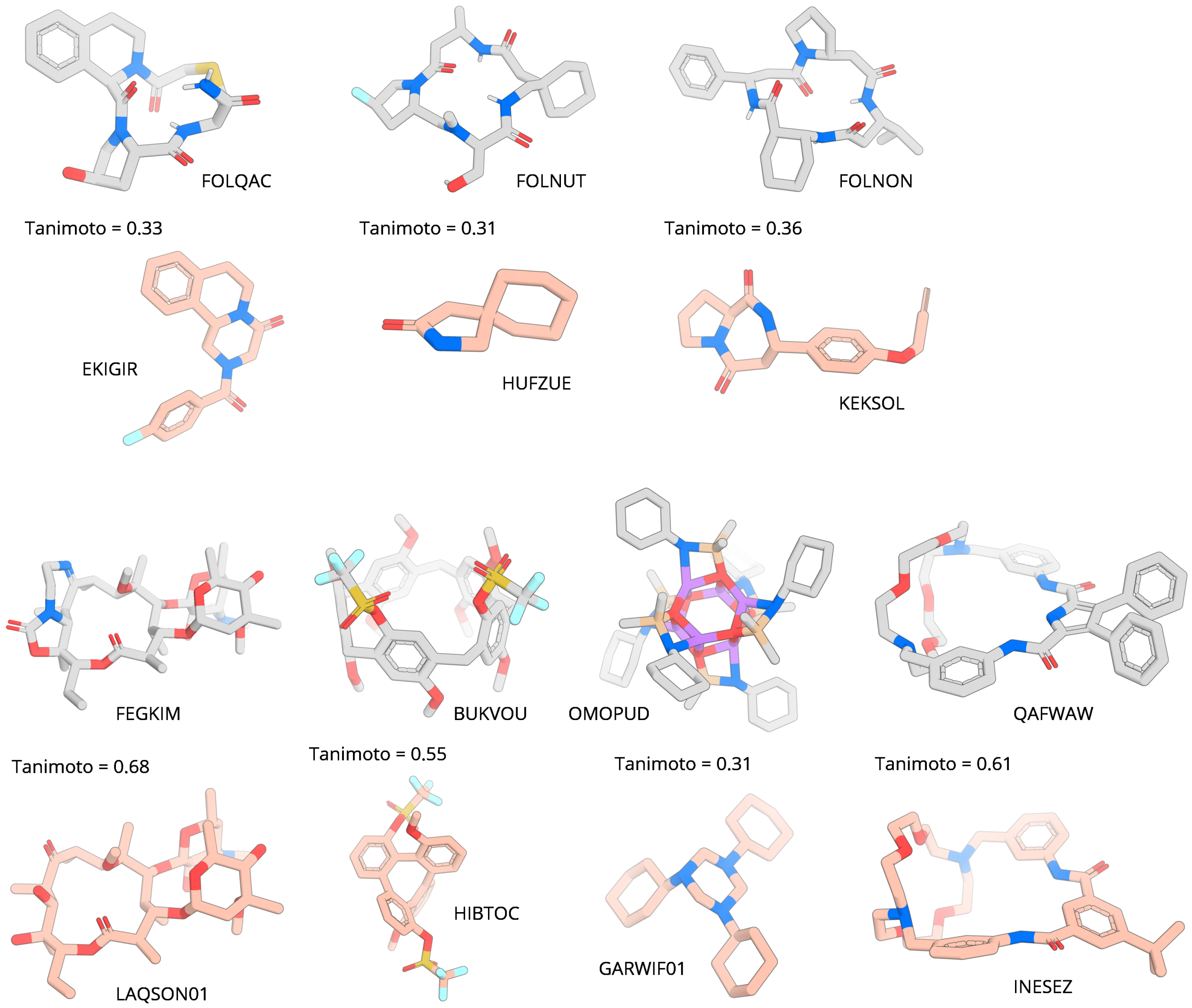


**Figure S3**. **Closest examples from the training set for the test molecules shown on Fig. 2D**. The former are shown in pale red while the latter are in grey. Respective CSD IDs are shown next to each molecule.

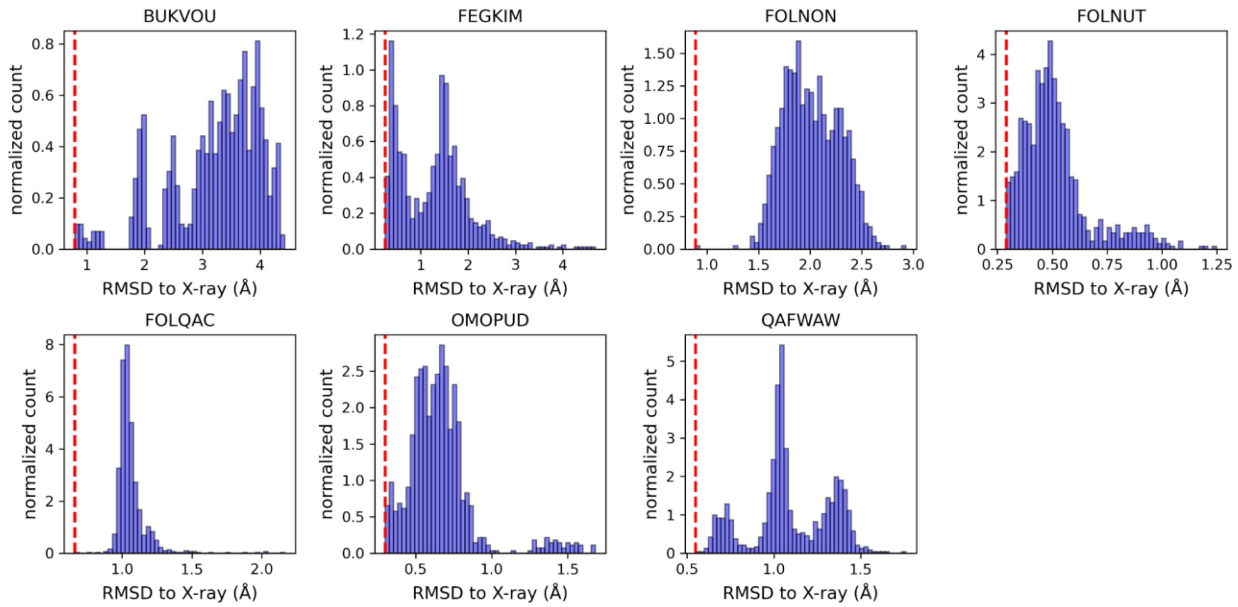


**Figure S4**. **RMSD distribution for all PLACER models generated for molecules shown on Fig. 2D.** Red dashed lines correspond to conformers with the lowest RMSD.

#### 2. Astex non-native benchmark

To evaluate PLACER's performance in ligand docking tasks we applied it on the Astex non-native benchmark set, consisting of 65 target molecules, placed into their non-native protein structures, with a total of 1112 unique ligand-protein pairs.(2) This means, if protein X was crystallized with molecules $M_1$ (backbone $X_1$) and $M_2$ (backbone $X_2$) in the same pocket then in this benchmark we evaluate the docking of molecule $M_1$ into backbone $X_2$. The benchmark dataset was prepared by aligning backbone $X_1$ to $X_2$, and copying the coordinates of $M_1$ into backbone $X_2$, deleting molecule $M_2$ and any other ligands. PLACER was run on each protein-ligand pair, generating 1000 models. The performance of PLACER was evaluated by ranking the 1000 models based on three confidence metrics (pRMSD, pLDDT, pLDDT-pDE), and using RMSD of the highest confidence model as the result.

An example PLACER run command was as follows:

```
run_PLACER.py --ifile input.pdb -n 1000 --rerank prmsd
```

This will generate two outputs: `input.csv` containing the scores (pRMSD, plDDT, plDDT-pDE) and RMSD values of each model, ranked by pRMSD from lowest (best) to highest, and `input_models.pdb` containing the geometries of each of the 1000 models. The b-factor column in the PDB file represents the pRMSD metric for each atom. pRMSD value in the CSV file is averaged over all atoms of only the ligand.

We further explored the impact of applying physics-based minimization on the PLACER-produced models. For this, we used the minimization protocol available in GALigandDock(9) on all of the produced 1000 models. This protocol was applied using a Rosetta XML input (`eval.xml`) and the following run command:

```
$ROSETTA/main/source/bin/rosetta_scripts -s {input.pdb} @flags.hb -parser:protocol eval.xml -extra_res_fa {LIG.params} -constrain_relax_to_start_coords -gen_potential -overwrite -beta_cart -no_autogen_cart_improper -out:levels all:300 protocols.ligand_docking.GALigandDock:300 -out:prefix {outdir}/ -out:file:scorefile {outdir}/scores.sc -crystal_refine -mute core basic -constant_seed -jran 1234567
```

The produced models were ranked based on the computed dG metric, and the RMSD of the best model used as the final result.

**eval.xml**

```
<ROSETTASCRIPTS>
  <SCOREFXNS>
    <ScoreFunction name="genpot_soft" weights="beta_cart">
      <Reweight scoretype="fa_rep" weight="0.2"/>
    </ScoreFunction>
    <ScoreFunction name="genpot" weights="beta_cart">
      <Reweight scoretype="coordinate_constraint" weight="0.2"/>
    </ScoreFunction>
  </SCOREFXNS>

  <TASKOPERATIONS>
  </TASKOPERATIONS>

  <FILTERS>
  </FILTERS>

  <MOVERS>
    <GALigandDock name="dock" scorefxn="genpot_soft" scorefxn_relax="genpot"
                  runmode="eval" turnon_flexscs_at_relax="1" contact_distance="8.0" >
    </GALigandDock>
```

```
  </MOVERS>

  <PROTOCOLS>
    <Add mover="dock"/>
  </PROTOCOLS>
  <OUTPUT scorefxn="genpot"/>
</ROSETTASCRIPTS>
```

**flags.hb**

```
-beta
-score::hb_don_strength hbdon_GENERIC_SC:1.45
-score::hb_acc_strength hbacc_GENERIC_SP2SC:1.19
-score::hb_acc_strength hbacc_GENERIC_SP3SC:1.19
-score::hb_acc_strength hbacc_GENERIC_RINGSC:1.19
-no_autogen_cart_improper
```

**Table S3. Astex non-native benchmark results with and without ligands at TC = 1.0 and TC ≥ 0.5 between the test set and the PDB training set.**

| | *Full Astex non-native set* | | *Omitted TC = 1.0** | | *Omitted* ***TC ≥ 0.5**** | |
|---|---|---|---|---|---|---|
| *Metric* | *< 1* Å **accuracy** | *< 2* Å **accuracy** | *< 1* Å **accuracy** | *< 2* Å **accuracy** | *< 1* Å **accuracy** | *< 2* Å **accuracy** |
| *pRMSD* | *41.8%* | *82.4%* | *40.8%* | *82.0%* | *39.2%* | *83.1%* |
| *pLDDT-1D* | *39.9%* | *78.6%* | *39.3%* | *78.3%* | *38.6%* | *80.2%* |
| *pLDDT-2D* | *37.6%* | *77.4%* | *36.5%* | *77.0%* | *34.5%* | *77.0%* |

** 22 examples with TC = 1.0 ligands KAI, SWA and VIB were omitted from the benchmark set*

*** 1296 examples with TC ≥ 0.5 ligands PH2, 675, AZM, A3M, SWA, NLA, IAD, BFL, IXM, STL, P16, 097, 3AR, GEO, THM, TQ3, NCT, VIB, AD3, KAI, SOX were omitted from the benchmark set.*

*Table S4. Tanimoto similarity coefficients for each of the ligands in the Astex non-native test set, compared against the PLACER PDB training set.*

| Astex ligand | Highest Tanimoto | Training ligand with highest Tanimoto | | Astex ligand | Highest Tanimoto | Training ligand with highest Tanimoto |
|---|---|---|---|---|---|---|
| PH2 | 0.564 | 2PH | | PM2 | 0.359 | 4PI |
| RQ3 | 0.471 | E0O | | SCT | 0.371 | AZZ |
| ID5 | 0.325 | IFB | | PFA | 0.346 | 64L |
| JE2 | 0.476 | E14 | | BYS | 0.455 | HF2 |
| MTI | 0.426 | IRP | | P16 | 0.608 | 96M |
| AO5 | 0.385 | JOV | | 097 | 0.545 | WR2 |
| 675 | 0.623 | J4X | | MOA | 0.415 | KP3 |
| CIA | 0.297 | L8X | | PFP | 0.378 | LIF |
| AZM | 0.619 | QOG | | MC9 | 0.210 | 4D8 |
| E4D | 0.421 | RAL | | 3AR | 0.619 | ARG |
| NDR | 0.492 | 3WF | | GIO | 0.388 | CHQ |
| A3M | 0.710 | WKM | | GEO | 0.593 | CTN |
| BIT | 0.303 | YCJ | | 984 | 0.317 | 59U |
| **SWA** | **1.000** | **SWA** | | LS1 | 0.382 | QYZ |
| BDI | 0.341 | OHJ | | TQD | 0.262 | W0S |
| STC | 0.419 | VWS | | FR4 | 0.351 | N0E |
| BSM | 0.374 | EH3 | | THM | 0.667 | AZZ |
| TNK | 0.362 | 4WF | | PH7 | 0.339 | LJ5 |
| CMU | 0.245 | I3V | | PAF | 0.422 | TXI |
| NLA | 0.541 | NOA | | BCZ | 0.349 | 3XR |
| LI9 | 0.409 | JQP | | PVB | 0.447 | C1W |
| SKF | 0.429 | 2CK | | TQ3 | 0.744 | JZM |
| IAD | 0.556 | E9M | | NCT | 0.575 | HNK |
| BFL | 0.564 | 9KL | | **VIB** | **1.000** | **VIB** |
| IBA | 0.313 | 4RB | | CEL | 0.467 | 4J8 |
| FSN | 0.384 | UIR | | 905 | 0.330 | J2Q |
| IXM | 0.528 | FEF | | AD3 | 0.527 | RFZ |
| BAU | 0.351 | 4WF | | CRZ | 0.477 | 6H6 |
| STL | 0.543 | R3S | | **KAI** | **1.000** | **KAI** |
| HUP | 0.226 | OCH | | CMB | 0.488 | IMA |
| BNE | 0.245 | JAN | | SOX | 0.629 | PNN |
| BIR | 0.477 | BEY | | 198 | 0.389 | 4CX |
| ROF | 0.338 | W3B | | | | |

### 3. Computational analysis of RA95 series of designed and evolved retroaldolases

To evaluate PLACER's ability to generate ensembles of enzyme active sites and use these models to discriminate between lower and higher activity variants we conducted an analysis on an existing set of enzymes with known activities. We chose the RA95-series of retroaldolase enzymes because there are high resolution crystal structures and detailed kinetic parameters of enzymatic activity for multiple variants along the evolutionary trajectory. The active sites of these enzymes center around nucleophilic lysine (K210 in RA95.0 and K83 in RA95.5 onwards) supported by an increasingly more sophisticated network of polar residues hydrogen bonding to the lysine and the substrate. In the reaction the lysine performs a nucleophilic attack at the carbonyl group of a beta-hydroxyketone substrate, which after shuffling of protons and departure of a water molecule undergoes a C-C bond cleavage reaction, yielding ultimately acetone and fluorescent 6-methoxy-2-naphthaldehyde (MeNA) products in case of the model methodol substrate (Fig. 4A). To perform the PLACER analysis on these active sites we focused on the covalently modified lysine at the different stages of the reaction - apo, methodol-carbinolamine (step 1), methodol-imine (step 2), acetone-imine (step 3), acetone-carbinolamine (step 4). We considered four different enantiomers of the methodol-conjugate; enantiomers with the lowest average predicted RMSD were used for analysis across the protein panel. To avoid irregularities of the crystal structures (missing loop densities, alternative atom placements), we collected sequences of the RA95 retroaldolases and predicted their structures using ColabFold following relaxation of the best ranking model with AmberRelax(10). Resulting models had Cα RMSDs between 0.15 and 0.47Å to corresponding crystal structures and were used for PLACER computations. We generated 50 samples with PLACER at each step and found that both the RMSD and the predicted RMSD (pRMSD) of the conjugated methodol correlated well with the experimentally determined catalytic activity of these variants (Fig. S2 and 4B). It is noteworthy that the trend is stronger when focusing on the methodol-conjugated lysine, and non-existent for acetone-conjugated lysine, highlighting PLACER's ability to consider the impact of the true substrate.

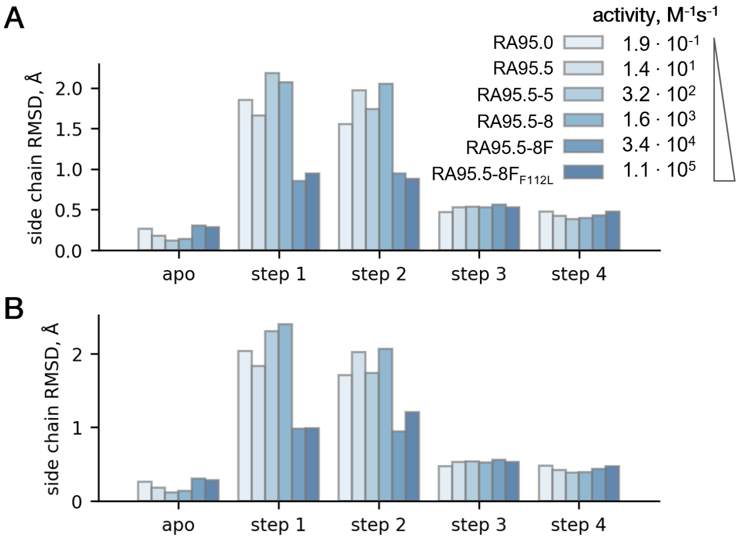


**Figure S5.** Five intermediate states of the RA reaction with methodol were modeled by PLACER for the RA95 series of designs to probe preorganization of the active site lysine and its modifications. Shown is the analysis based on the RMSD of the free lysine (*apo*) or conjugated lysine atoms except N, Cα, C, O atoms. **A)** Enantiomer with lowest RMSD is used for step 1 and step 2 in the top panel. **B)** RMSDs for *R,R*-enantiomer are in the lower panel.

### 4. Computational de novo design of retroaldolases in NTF2 fold

We explored the ability of PLACER to aid in the selection of higher activity de novo designed enzymes, using the retro-aldol reaction as an example.

A short summary of the workflow for evaluating PLACER in the context of enzyme design:

1. Design methodology development using de novo generated NTF2 scaffolds
2. Experimental identification of most viable active site configuration
3. Development of PLACER "selection model" based on the data from step 2
4. IVTT based increased throughput retroaldolase activity testing assay
5. Individual purification and activity testing of selected designs
6. In depth PLACER analysis of the experimentally tested designs

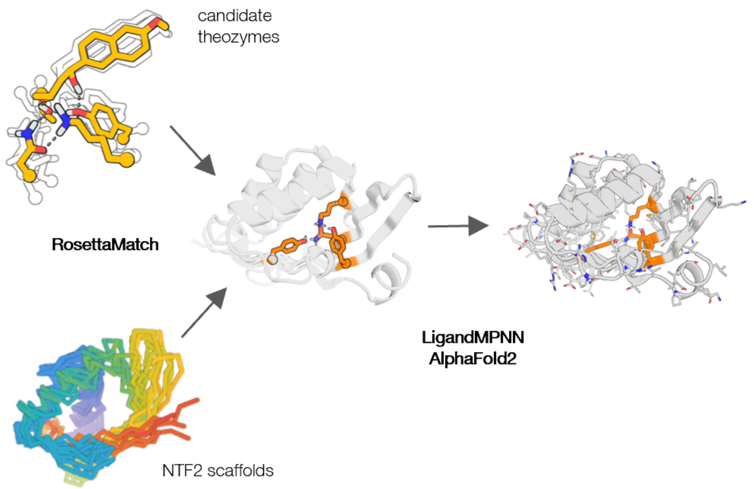


**Figure S6.** Overview of the design workflow.

Successful design process requires to address following challenges: generation of large number of designable backbone variations of NTF2 fold, identification of potential locations where catalytic quartet can be installed and finally coming up with selection criteria to downsample number of designs for experimental testing. To address the first challenge we used RoseTTAFold joint inpainting (RFjoint2)(11) to generate complete NTF2 scaffolds by randomly retaining 3 or 5 contiguous residues in major strands and helices of the randomly picked "seed" NTF2 structure, and allowing RFjoint2 to fill in missing pieces of the backbone. For each generated scaffold TM score relative to the "seed" structure was computed and 10000 scaffolds with the score greater than 0.62, which ensured NTF2-like topology, were selected for the next stages.

To install the catalytic quartet we used a two step procedure. We first used a modified XML-Matcher protocol(12) to determine what positions in the NTF2 fold might be broadly compatible with geometric requirements of the four residues interacting with the ligand and forming a hydrogen bond network. Instead of exhaustive enumeration of all possible spatial arrangements consistent with desired connectivity, we implemented a guided Monte-Carlo docking procedure to bring disjointed randomly placed residues together to form a theozyme as illustrated in Figure S4. By repeating docking trajectories of the disembodied sidechain functional groups with each of the 4 possible stereoisomers of the carbinolamine intermediate, and removing outputs that did not have desired connectivity, we generated 4 ensembles of several thousands structures each. The number of successful trajectories varied substantially between stereoisomers (*R,R* – 4352 out of 5000, *R,S* – 2177, *S,R* – 4373, *S,S* – 2910), suggesting structural preferences of the catalytic quartet. After grouping generated structures by similarity of *nucleophile* (“nuc”) and *shuttle* (“sht”) positions (183 total clusters generated, split between *R,R* – 20, *R,S* – 23, *S,R* – 77, *S,S* – 63), geometric parameters describing rigid body transformations between reference atoms of the ligand and reference atoms of the protein side chains of cluster members were extracted and written into XML-formatted constraint files. Additional theozyme diversity was incorporated by allowing the *bridge* (“brd”) residue to be either Asn or Gln, and *support* (“sup”) to be Tyr, Ser or Thr, resulting in 6 possible theozyme compositions. This approach is not as thorough in exploring all possible ways to arrange theozyme components in space as it would be with HBNetGen(12), but more rapidly generates diversified theozyme coordinates and enables exploration of alternative site descriptions. This more rapid feedback was necessary as initial attempts to match catalytic quartet theozymes in native scaffolds resulted in a large fraction of the output having theozyme residues placed on the unstructured looped regions of the scaffolds. To enrich sampled theozyme geometries with those compatible with catalytic residues projecting from ordered secondary structure elements, we repeated the docking protocol with theozyme residues in tripeptide segments of helical or extended backbones; this resulted in more matches in regular secondary structure elements. In total 6*183=1098 individual constraint files were generated and used for matching.

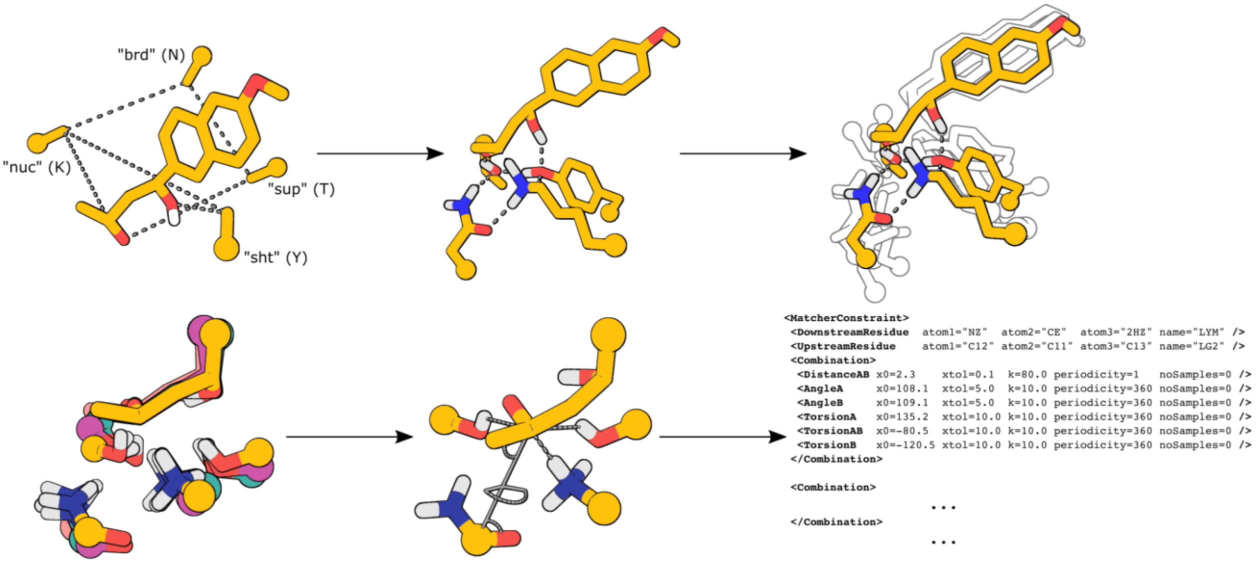


**Figure S7.** Preparation of theozyme geometries with a guided Monte-Carlo docking procedure and converting them into xmlMatcher formatted constraints. (top panel) each docking trajectory is initialized form theozyme components randomly oriented in space and brought into desired geometry by cycles of rigid body perturbations and restrain guided minimization with Rosetta; (bottom panel) rigid body orientation geometries of chemical motifs are extracted from ensemble and written into xmlMatcher formatted text file

Since the protocol is computationally intensive, we performed the computation for 500 out of 10000 randomly selected NTF2 backbones. We identified several combinations of positions and theozyme residue types that frequently produced reasonably good approximations of the desired geometry in multiple backbones. Following this preliminary stage, we used structurally equivalent positions in all 10000 scaffolds to install theozyme residues and design the rest of the positions in the backbone using structure conditioned sequence design in the presence of the substrate (LigandMPNN)(13) to generate multiple sequences for each backbone. AlphaFold2(7) was used to predict the structure of each sequence (in single sequence prediction mode, using model 4 ptm and 3 recycles), and models with lDDT higher than 85 created a pool of designs subjected to various *in silico* selection strategies.

#### 4.1. Identification of focus theozyme configuration

The space of possible theozyme configurations (catalytic residue placement and amino acid identities) in any given protein topology is finite and limited by the geometry of the interacting theozyme side chains. However, not every configuration, while consistent from geometric perspective, is going to be inherently catalytically competent. Therefore, to be able to focus on assessing the role PLACER in the design workflow, we needed to establish conditions such that a reasonable number of active designs could be expected in the batch of experimentally tested variants.

We ran a preliminary experiment where a batch of 90 designs was split between three most promising theozyme configurations (Table S3) identified using XML-Matcher protocol (34, 35, and 21 designs for configurations 1, 2 and 3 respectively).

**Table S5. Tested configurations of and placements of retro-aldolase theozymes**

| Configuration | "nuc" | "brd" | "sup" | "sht" |
|---|---|---|---|---|
| 1 | K80 | N78 | S91 | Y108 |
| 2 | K62 | Q78 | Y104 | Y89 |
| 3 | K106 | Q104 | Y13 | Y52 |

Selection criteria for individual designs were based on how well side chains of the theozyme residues recapitulated desired geometry in the AF2 predicted structures with pLDDT > 85 and did not include PLACER metrics. Results of activity testing using IVTT protocol (Figure S5) suggested that one of the theozyme configurations may be more designable using our computational pipeline.

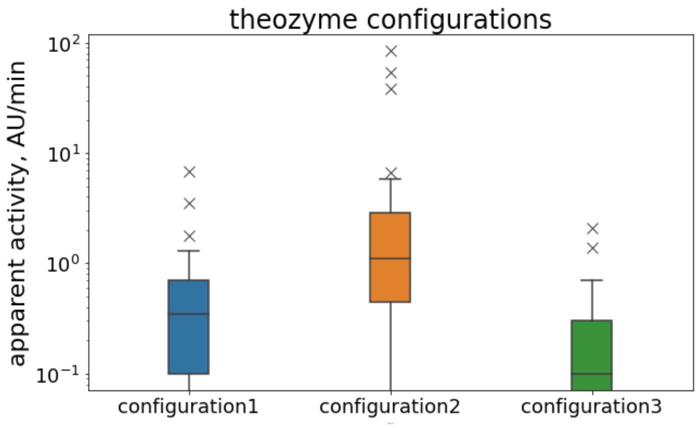


**Figure S8.** Distribution of apparent activities for designs with various theozyme configurations. Configuration 2 appears to be more compatible with underlying requirements of retro-aldolase active site geometry and was used to generate a pool of designs for testing PLACER selection strategy.

As a result of this experiment we decided to use configuration 2 to generate the common pool of designs used to test computational selection strategies.

#### 4.2. Application of PLACER for design selection

To test various downsampling selection strategies we generated a common pool of designs by installing theozyme configuration 2 into 10000 NTF2-like scaffolds (368 scaffolds in which at least one component of the theozyme could not be mapped into structurally equivalent environment were omitted from design calculations). For each of the 9632 scaffolds 8 sequences were generated with LigandMPNN, keeping residue type of the theozyme positions fixed, and for each sequence its structure was predicted with AlphaFold2. Structures with pLDDT ≥ 85 ( 17180 in total ) formed the pool of designs used to select variants for experimental testing.

(i) First, "naive" strategy relies on the hypothesis that AlphaFold2 predicted positions of the theozyme side chains in the *apo* form of the design correlate with actual conformations of these residues both in *apo* and *holo* form of the design. From the pool of designs we would select ones where the AlphaFold2-predicted *apo* structures have the theozyme residues participating in required hydrogen bonds.

(ii) Second strategy relied on using PLACER as the design evaluation tool. For each design, we would run PLACER 50 times, creating an ensemble of conformations. A set of six metrics, such as uncertainties in the positions of the polar atoms, and average distances between hydrogen bonding atoms of the theozyme residues were computed across ensemble members.

Base selection strategy, dubbed "naive", used a set of interatomic distances between polar atoms of the theozyme side chains "nuc(NZ)"-"brd(OE1)", "sup(OH)"-"brd(NE2)", "nuc(NZ)"-"sht(OH)" in the predicted structure as a proxy for formation of the hydrogen bonding network in the theozyme in the absence of the ligand. This was complemented by RMSD of the theozyme side chain heavy atoms between AlphaFold2 predicted structure and a structure of the pre-LigandMPNN design with theozyme geometry optimized to reflect preconceived vision of transition state stabilization by the protein. Numerical values for the distances and RMSD cutoffs were adjusted to result in 90 designs passing the selection ( RMSD < 1.73 Å, "nuc(NZ)"-"brd(OE1)", "sup(OH)"-"brd(NE2)" and "nuc(NZ)"-"sht(OH)" between 2.0 and 3.3 Å). DNA sequences encoding for these designs were obtained as IDT eBlocks™ and their activity tested following the IVTT protocol.

Selection strategy, dubbed "PLACER", utilizing PLACER to assist in picking the most promising designs from the same pool of 17810 variants relies on a set of geometrical and uncertainty metrics from generating an ensemble of 50 structures with PLACER, predicting interactions between the ligand and the active site residues in the context of AlphaFold2 predicted backbones. The obtained metrics included distances between polar atoms of the theozyme side chains and uncertainty (*pRMSD*) in polar atoms positions computed over the ensemble of 50 models. To identify a smaller subset of PLACER metrics we trained a simple logistic regression model to classify designs as active or inactive on a set of designs characterized in the preliminary experiment performed to identify focus theozyme configuration.

#### 4.3. IVTT based retroaldolase activity testing assay

We developed a protocol for medium throughput semi-quantitative retro-aldolase activity testing of the designs using in vitro transcription translation system (IVTT). We obtained linear duplex DNA, encoding protein of interest, flanked by appropriately spaced T7 promoter and T7 terminator sequences as eBlocks™ gene fragments from IDT, and by mixing it directly with 2 µL components of PurExpress IVTT system produced enough protein to screen for active variants. Genes of several control proteins with a range of activities previously determined in purified form were included with each batch of tested designs to assist in ranking activity of the new variants, and help to compare experiments run under different experimental conditions. It should be noted that in this format of the experiment, concentration of the soluble protein remains unknown and therefore activity level should be treated as apparent activity defined as a product of the amount of soluble protein and intrinsic activity of each variant. It is possible to make assay more quantitative by fusing protein of interest to fluorescent protein reporter at the expense of decreased amount of retro-aldolase catalyst synthesized. Typically, a batch of 90 designs and 6 controls were used to evaluate performance of a particular computational workflow.

#### 4.4. Characterization of individual variants

To get more accurate and quantitative data describing catalytic activities of the designs and validate IVTT based assay, we cloned, expressed in Escherichia coli, purified and determined Michaelis-Menten parameters for twelve designs with highest apparent activity from "naive" or "PLACER" group (24 proteins in total). Additionally, we purified and tested under similar experimental conditions three previously characterized proteins possessing retro-aldolase activity. This group of control proteins consists of published RA95.5-8F(14), RA110-4.6(15), and RA**β**b-16.2(16) retro-aldolases computationally designed and subjected to various extent of directed evolution optimization, leading to significant improvements in catalytic proficiency.

Under generic unoptimized conditions, soluble expression levels in *E. coli* varied almost 100-fold among 24 proteins, indicating the possibility that when uncorrected for concentration, IVTT determined activities may be unreliable in recovering high activity variants. However, activity of the designs expressed in terms of $k_{cat}/K_M$ correlate qualitatively ($R^2$ ~0.4) with apparent activity measured in IVTT experiment, suggesting that findings on the smaller subset of purified proteins can be, to certain extent, extrapolated onto the larger subset of proteins characterized in IVTT format. Moreover, the highest activity variants in the "PLACER" group found in IVTT experiment are indeed the highest activity variants by their $k_{cat}/K_M$ values, except two variants in IVTT assay that appeared to have activity level beyond the dynamic range of the assay, but in purified form demonstrated modest levels of activity. Mean $k_{cat}/K_M$ values in "naive", "PLACER", and "control" groups are 250, 2300, and 80,000 $M^{-1}$ $min^{-1}$ respectively.

Furthermore, results obtained with purified proteins confirm the initial observation that designs cnRA-34 and cnRA-50 in the "PLACER" group, with $k_{cat}/K_M$ values of 6900 and 11,000 $M^{-1}$ $min^{-1}$ respectively, are the most efficient computationally designed catalysts that were not subjected to additional experimental optimization, and being similar to the activity of the RA**β**b-16.2 (12,000 $M^{-1}$ $min^{-1}$, 5 rounds of evolution) and only surpassed by RA95.5-8F (240,000 $M^{-1}$ $min^{-1}$, 18 rounds of evolution; it should be noted that published estimates for $k_{cat}/K_M$ of this enzyme are 5-10 times higher most likely due to phosphate based buffer used in this study, which was shown to have strong inhibitory effect on retro-aldolases built on 1a53 scaffold(17)) (Figure S6).

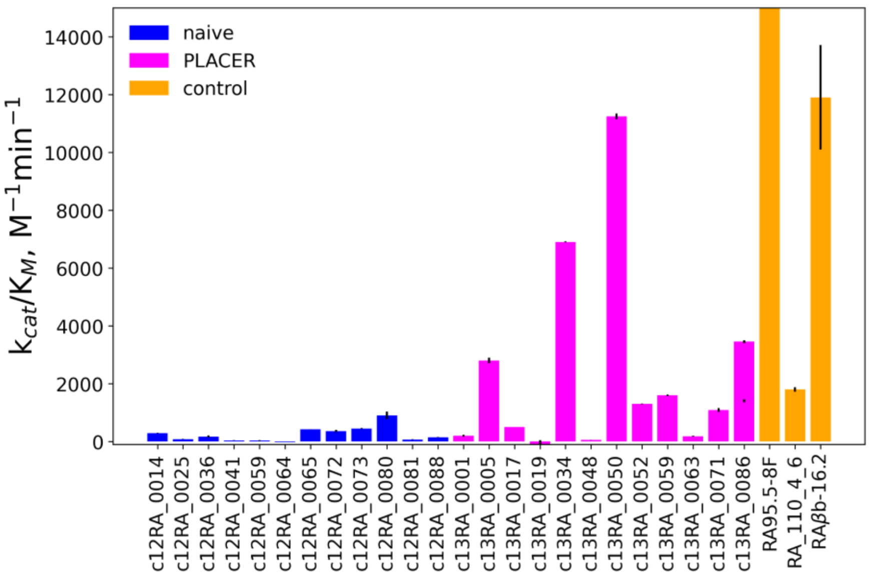


**Figure S9.** Values of $k_{cat}/K_M$ for top 12 designs from "naive" and "PLACER" group.

#### 4.5. In depth PLACER analysis of the experimentally tested designs

Obtaining a substantial number of designs with detectable level of activity allowed us to perform analysis analogous to that of the RA95 series of retroaldolases (Fig. 4B and Figure S2) on all experimentally characterized (either in IVTT or in purified protein format) de novo NTF2 designs (548 total), and see the level of discrimination between active and apparently inactive variants according to PLACER ensembles. For each design we generated 50 samples for each enantiomer of methodol-carbinolamine (step 1) or methodol-imine (step 2) intermediates. Due to an opening created by PLACER pose cropping (<600 atom crop) of NTF2 topology, some models in the PLACER ensemble have an incomplete representation of the pocket and allow the buried catalytic residues to "escape" the protein active site, artificially increasing values of ensemble RMSD. To mitigate this effect, we removed from the analysis members of the ensemble deviating more than two standard deviations from the ensemble centroid. We found that both the RMSD and the pRMSD of methodol atoms in conjugated lysine correlated well with the experimentally determined catalytic activity of these variants (Fig. S7 and 4F). It should be noted that none-zero fraction of designs may be falsely labeled as inactive in IVTT assay due to errors in coding DNA eBlock synthesis or low soluble expression in a non-cellular environment, and may contribute to distribution overlap, making discrimination less clear.

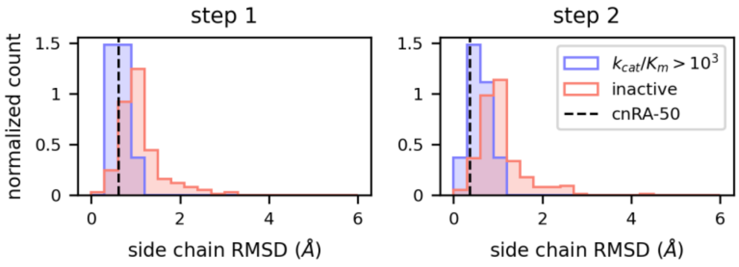


**Figure S10.** PLACER active site pre-organization ensemble metrics (average RMSD of conjugated lysine atoms except N, Cα, C, O to ensemble centroid positions) in steps 1 and 2 enrich for selecting more active de novo designed retro-aldolase enzymes.

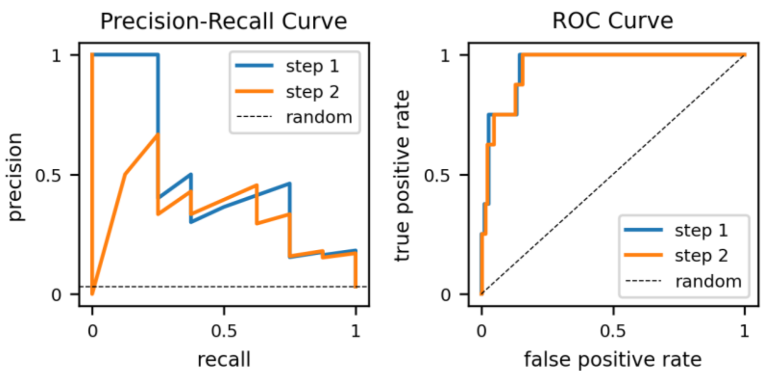


**Figure S11.** Separation between pRMSD distributions for active and inactive designs from Fig. 4F shown in terms of PR and ROC curves.
